# Heterospecific developmental models highlight the role of *FBXO32* in neural crest lineages and the significance of its variant in pediatric melanoma

**DOI:** 10.1101/2023.03.10.532100

**Authors:** Vargas Alexandra, Guillermain Clémence, Gorojankina Tatiana, Creuzet Sophie

## Abstract

Melanoma is a rare sporadic pediatric cancer caused by *de novo* genetic accidents. To analyze the involvement of *FBXO32* of unknown biological significance, we set up original developmental in *vivo*, *ex vivo*, and *in vitro* models to unmask the biological role of *FBXO32* in cephalic and trunk neural crest lineages. We show that *FBXO32* silencing impairs melanocyte proliferation, differentiation, and vascularization. In rescue experiments, co-electroporation with the human wild-type gene compensates for the silencing of the endogenous gene and restores normal melanocyte phenotypes. By contrast, co-electroporation with the human variant, *FBXO32^G139S^*, alters melanocyte differentiation, pigment synthesis, triggers migration to the dermis, and favors angiotropism. Transcriptomic analysis reveals targets associated with melanocyte transformation, cytoskeleton remodeling, and focal adhesion. We show that *FBXO32* orchestrates unexpected interactions with *ASIP* for melanogenesis and *BAP1* for the post-translational ubiquitination system. We propose *FBXO32* as a tumor suppressor gene capable of preventing the onset of pediatric melanoma.

## INTRODUCTION

Melanoma is a skin cancer of transformed melanocytes, pigment-producing cells derived from the embryonic Neural Crest (NC). Malignant cutaneous melanoma is a rare neoplasm in the pediatric age group, accounting for 4% of all cancers in children aged 15 to 19 ^1^. The incidence in children is stable at <1 per 100 000 person/year, whereas adolescent melanoma has increased over the past 20 years related to unprotected sun exposure ^2^. Childhood melanoma risk factors include many nevi or giant congenital melanocytic nevi (CMN) and dysplastic nevus syndrome. Other risks are associated with predisposition syndromes such as *Xeroderma pigmentosum*, immunosuppression, and predisposition to a genetic disorder ^3,4^.

Childhood melanoma’s genetic landscape differs from adult counterparts ^5–7^. The germline *CDKN2A* mutation identified at 4-6% in adult melanoma is not significantly associated with children with an atypical nevus and familial history ^6–9^. While adult melanoma creates a permissive condition for developing malignant melanoma upon UV exposure, childhood melanoma arising from CMN demonstrates a lower frequency of UV-related mutations ^10,11^. Furthermore, *NRAS^Q61R^,* prone to drive 70% to 95% of *in-utero* CMN, indicates that early-onset childhood melanoma development does not require UV exposure ^12^. Similarly, in CMN, the *BRAF^V600F^* low-frequency mutation develops *in-utero,* and melanoma arises in the non-exposure area. The *BRAF* chromosomal translocations are involved in CMN activation of melanocytic tumors ^12,13^. Although the melanocortin-1 receptor (MC1R) gene is highly polymorphic and associated with two alleles variants, *R* and *r,* in red hair color phenotype, children and adolescent melanoma have a higher prevalence of *MC1R r* variants than in adult melanoma through a pigmentation-independent pathway ^14,15^. This genetic evidence in childhood melanoma suggests the involvement of biological pathways other than pigmentation through UV sensitivity. In conventional pediatric melanoma (CM) dysregulation of the *PTEN* tumor suppressor gene, the negative regulator of the phosphatidylinositol-3 kinase/AKT pathway is associated with *BRAF^V600F^* mutation ^6^. In contrast, *NRAS* mutations do not require the cooperation of *PTEN* mutations in CMN ^6^. The somatic mutation in the *TERT* promoter disrupts the telomeric DNA length and chromosomal stability frequently observed in pediatric CM and adolescents ^17–19^. The germline *TP53* mutation is an initial malignant tumor in a pediatric melanoma associated with Li-Fraumeni syndrome ^18^. Moreover, loss of the BRAC1-associated protein 1 (*BAP1)* expression coincides with schizoid-appearing benign and malignant melanocytic proliferations in pediatric melanoma ^19,20^. Besides, early-onset cancers are a classic sign of a genetic predisposition, but most pediatric melanomas are clinically sporadic, without correlation to any inherited familial genetic history ^3^.

Compelling evidence pointed to the critical role of ubiquitination and deubiquitination in metabolic reprogramming in cancer cells ^21^. Among the ubiquitination pathway, the F-box proteins constitute a subunit of the ubiquitin protein ligase complex, SKP1–Cullin1–F-box protein E3 (SCFs). The F-box E3 proteins comprise three subclasses: the FBXW proteins containing WD-40 amino acid domains, FBXL proteins containing leucine-rich amino acid repeats, and FBXO, F-box-only proteins with uncharacterized domains ^22^. The F-box proteins recognize specific protein substrates for ubiquitination in a phosphorylation-dependent manner ^23^. Their functions involved cancer hallmark pathways such as cell cycle, DNA damage, epithelial-mesenchymal transition (EMT), and signaling pathways such as AKT/PI3K, AMPK/mTOR, AKT, NF-κB ^23,24^. According to published data, the gene *FBXO32*, localized at 8q24.13 in humans, has both tumor-suppressive and tumor-promoting roles in regulating proliferation and migration processes in different cancers. In breast cancer cells, its effect results in the *CTBP1* nuclear retention to control EMT and metastases ^25^. In melanoma, *FBXO32* was identified as a *MITF* target oncogene regulating cell migration and proliferation ^26^. The *FBXO32* dysregulation promotes EMT in urothelial carcinoma while acquiring platinum resistance ^27^. As a possible tumor suppressor, *FBXO32* transcription was epigenetically silenced through DNA methylation, disrupting TGF-β/SMAD4 signaling in ovarian cancer cells ^28–30^. In addition, overexpression of *FBXO32* promotes c-Myc oncogene degradation to mediate cell growth inhibition ^31^. Nonetheless, to date, the role of *FBXO32* in childhood melanoma is still elusive. Furthermore, the *FBXO32* gene degrades a zinc-finger transcription factor Krüppel-like factor-4 (KLF4) to suppress breast cancer progression via the p38 mitogen-activated protein kinase pathway ^32^.

Here, we explore the biological role of the *FBXO32* gene and test its variant’s significance for the occurrence of sporadic melanoma. The *FBXO32* mutation, c.415G>A (p.G139S), was identified in a 7-year-old melanoma patient without melanoma family history in blood-extracted DNA by constitutional exomes sequencing (200X) ^33^. We hypothesized that sporadic pediatric melanoma occurs as a *de novo* genetic “accident” in a parental gamete or post-zygotically during neural crest (NC) development. To understand the biological significance of the *FBXO32* gene in NC cell (NCC) development and the relevance of p.G139S mutation to melanomagenesis, we set up original developmental models to carry out longitudinal *in vivo, ex vivo* and *in vitro* functional analyses in the chick embryo. First, we tested the relevance of *FBXO32* for the normal development of NCCs at cranial and trunk neural crest levels (CNC and TNC, respectively). By RNAi-mediated silencing, we show that *FBXO32* silencing in CNC and TNC provokes aberrant melanocyte proliferation and differentiation. Then, we established a 3D skin explant model to investigate the role of *FBXO32* and p.G139S mutation during the specification of melanoblasts and their differentiation into melanocytes. We demonstrated that *FBXO32^G139S^* induces cytoskeleton remodeling through focal adhesion rearrangement, actin filaments disruption, and cell aggregate formation. Finally, RNAseq data mining determined gene expression profiles in CNC and TNC samples to identify the transcriptional signatures associated with *FBXO32^G139S^*. Our bioinformatic analysis unravels *FBXO32* as an upstream target of agouti signaling protein (ASIP) in the MC1R signaling pathway and *BAP1* deubiquitinase in post-transcriptional regulation. These data evidence the role of *FBXO32* as a tumor suppressor in melanocytic transformation at early developmental stages.

## RESULTS

### *FBXO32* expression in CNC and TNC cell populations

To evidence the expression pattern of *FBXO32* during NC and melanocyte lineage development, we performed whole-mount *in situ* hybridizations on chick embryos (Figure S1). At the early neurula stage, corresponding to the 4-somite stage (ss), *FBXO32* transcripts were detected at the margin of the neural plate (Figure S1A). At 24ss, *FBXO32* expression was observed in delaminating NCCs at the nasofrontal ectoderm, the retro-ocular region, and the third branchial arch (Figure S1B). The accumulation of FBXO32 transcripts was detected along the neural tube at 26ss (Figures S1D and S1E). Later in development, at 13 embryonic days (E), FBXO32 transcripts were detected in NCC-derived melanocytes differentiating in the epidermis (Figure S1F), while the dermis was devoid of *FBXO32* transcript and pigmented melanocytes. Immunostaining using anti-FBXO32 and anti-HNK1 antibodies (Abs) confirmed these observations. The HNK1 epitope is a marker of CNC and TNC during NCCs delamination and migration ^34^. Immunostaining with anti-FBXO32 and Phalloidin staining revealed FBXO32 accumulation in NCCs before and during their migration (Figures S1G, S1H, S1I, and S1J). Furthermore, at 24ss, FBXO32 immunostaining colocalized with anti-HNK1 positive delaminating CNC (Figures S1K and S1L). In the trunk, FBXO32 accumulates at the neural tube, and HNK1-positive TNC cells (Figure 1SM). These results demonstrate a specific *FBXO32* expression in CNC and TNC lineages.

**Figure 1:**
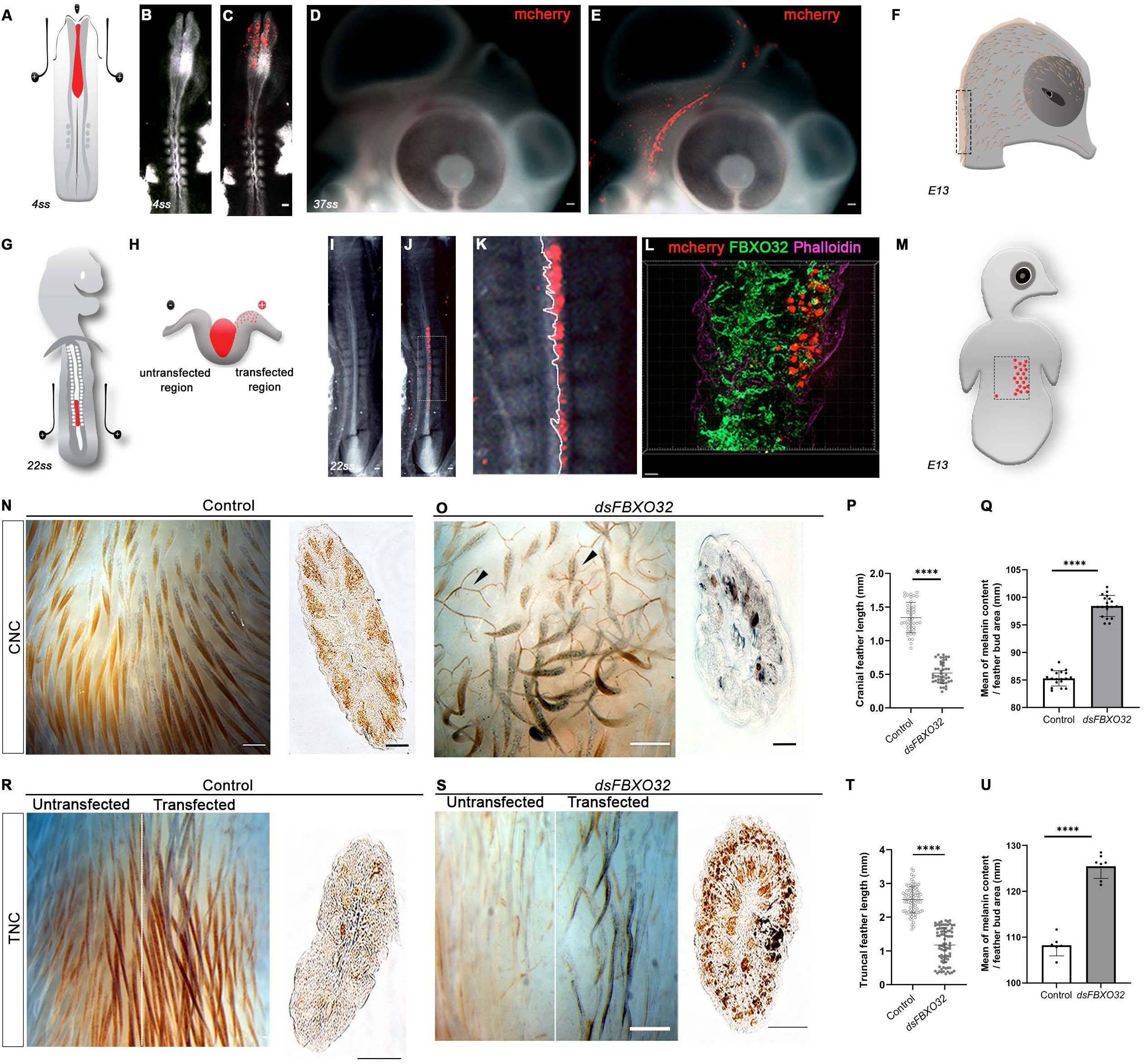
Silencing of *FBXO32* in CNC and TNC yields abnormal melanocyte differentiation in the head and the trunk. **A)** Experimental design for bilateral electroporation of CNC in 4ss chick embryo. **B)** Whole-mount embryo at 4ss, showing **C)** mCherry reporter gene accumulation in NCCs 3h after transfection **D)** Embryo’s head at 37ss where **E)** mCherry features NCC migration, two days after electroporation. **F)** Diagram of phenotype analysis at E13. **G)** Unilateral electroporation of TNC at 22ss. **H)** Diagram showing mCherry-positive transfected cells. **I)** Plasmid (red) injection at 22ss. **J)** Detection of mCherry-positive NCC delaminating after unilateral electroporation. **K)** Magnification of J. **L)** TNC expressing mCherry are immunolabelled with anti-FBXO32 mAb (green) 20h post-electroporation; Phalloidin (purple) staining outlines embryonic structures. **M)** Diagram of an E13 embryo where the dotted lines delineate the transfected region. **N)** In E13 control embryos, feather buds developing from transfected CNC exhibit pheomelanin accumulation (foxy pigmentation). On the transverse section (right), melanocytes fill the barb ridges in the epidermis. **O)** In *dsFBXO32-*transfected embryos, *FBXO32* silencing impairs melanogenesis and feather polarity. Arrowheads indicate the expanded vascularization into the dermis. On the transverse section (right), *dsFBXO32-* electroporated CNC forms aberrant melanocytes misoriented towards the feather’s dermal pulp. **P)** Box and whiskers plots showing the differences in feathers length (mm) between controls and *dsFBXO3*-treated embryos. **Q)** Graph representing the relative melanin content average per feather bud (mm) in the transfected area between control and *dsFBXO3-*treated embryos. **R)** In E13 control embryos, feather buds developing from transfected TNC exhibit normal foxy pigmentation and anterior-posterior polarity. The transverse section through the transfected area shows melanocyte localization and differentiation in feather buds. **S)** In embryos subjected to *FBXO32* silencing in TNC, melanocytes accumulate eumelanin in the transfected region. **T)** Box and whiskers plots highlight alterations in feather length *dsFBXO32*-electroporated embryos compared to control. **U)** Average melanin content per feather bud area (mm2) in control and *dsFBXO32*-treated embryos, summed from 4 control and 3 *dsFBXO32*-transfected embryos. Bars indicate the mean; error bars, SD; one-way ANOVA with Dunnett’s multiple comparison tests (\*\**P*<0.05, \*\*\*\**P*<0.0001, ns: not significant). Scale bars: 50µm.

### *FBXO32* silencing results in abnormal melanocyte differentiation in both CNC and TNC

To understand the role of *FBXO32* during NCCs development, we silenced the chicken *FBXO32* gene by delivering double-strand RNA molecules (dsRNA) designed against its cDNA (*dsFBXO32)*.

First, we performed *dsFBXO32* bilateral electroporation in CNC at 4ss in chick embryos (Figure 1A). The efficacity of NCCs bilateral transfection was monitored by co-electroporating with pCAGGS-mCherry (mCherry) plasmid driven by the cytomegalovirus (CMV) promoter (Figures 1B and 1C). Three days after CNC transfection, at 37ss, mCherry-positive NCCs populate retro-ocular and cranial regions (Figures 1D and 1E). The consequences of *FBXO32* silencing were analyzed 11 days post-electroporation in the cranial epidermis (Figure 1F). In E13 control embryos, feather development exhibited longitudinal growth following an anterior-posterior polarity and melanocytes with pheomelanin (foxy) pigmentation located on the basal lamina of barb ridges epidermis (Figure 1N). Following *FBXO32* silencing, we noticed abnormal feather polarity: skin morphogenesis was severely perturbed. Moreover, melanocytes exhibited pigment shift with massive eumelanin (black) accumulation and aberrant polarization towards the feather’s dermal pulp (Figure 1O). Quantifications of feather length and melanin content in *dsFBXO32* embryos showed that feathers were hypoplastic and harbored eumelanin accumulation (Figures 1P and 1Q). In addition, *FBXO32* silencing in CNC increased blood vessel density in the dermis (Figure S2A). Compared to controls, the vessel density in the cranial dermis was threefold after *dsFBXO32*-electroporation of CNC (Figure 2SB). Our results demonstrated the detrimental effects of *FBXO32* silencing in CNC on feather development, pigmentation synthesis, and vessel development.

**Figure 2:**
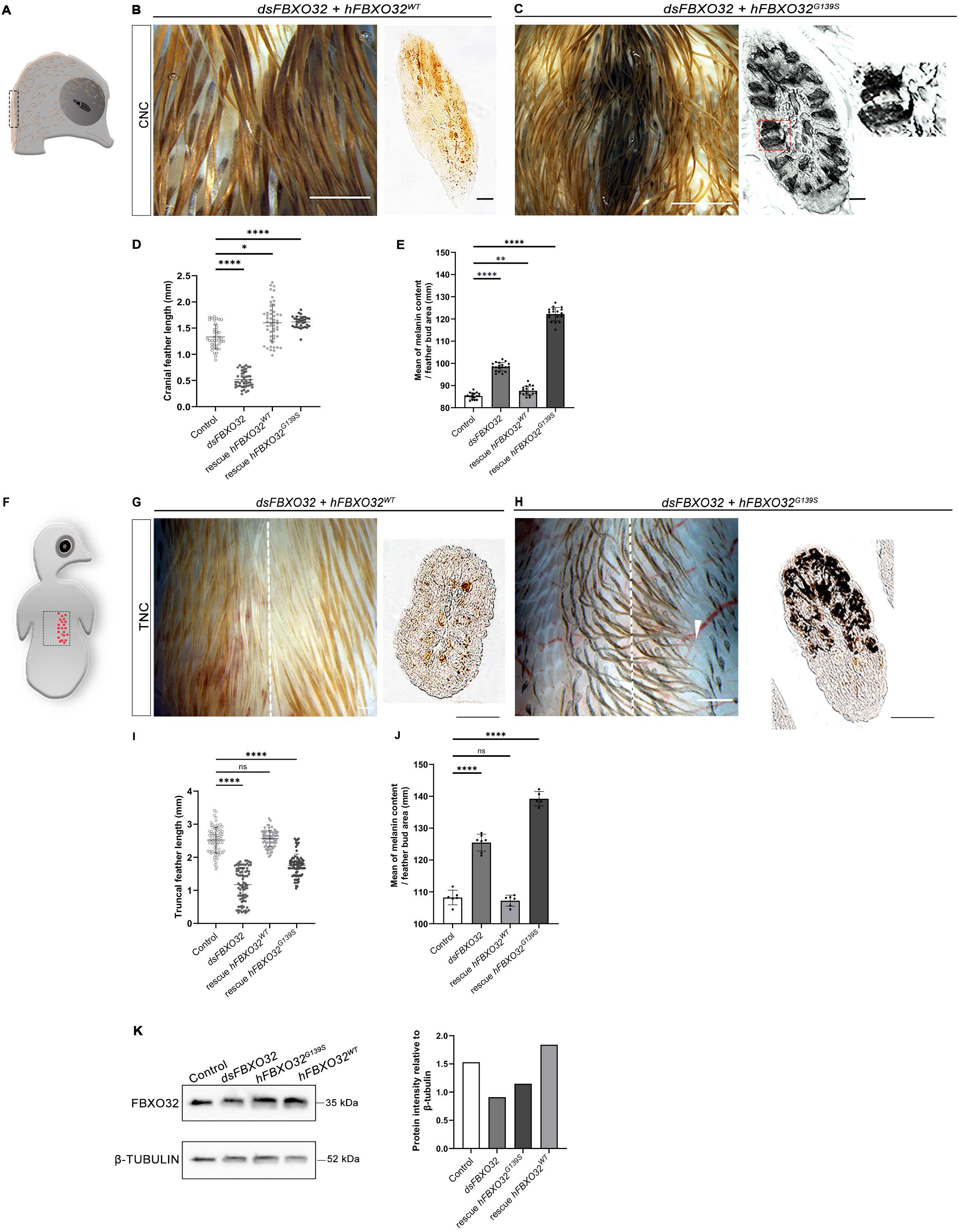
Human wild-type *hFBXO32^WT^*gene can restore the defects yielded by *FBXO32* silencing in NCC, but the *hFBXO32^G139S^* gene impairs melanocyte differentiation, feather morphology, and polarity. **A)** Diagram showing transfected cranial region. **B)** Dorsal views of E13 embryo subjected to *dsFBXO32*+*hFBXO32^WT^* transfection in CNC. On the feather bud section (right), melanocytes localize in barb ridges. **C)** Dorsal views of E13 embryo subjected to *dsFBXO32*+*hFBXO32^G139S^*transfection in CNC. On the feather bud section (right), melanocytes form dense and large eumelanin patches. Red squares show defective melanocytes’ polarity with filopodia oriented towards the dermis. **D)** Box and whiskers plots showing average feather length comparing the effects of *dsFBXO32*+*hFBXO32^WT^ and dsFBXO32*+*hFBXO32^G139S^*in CNC. **E)** Average melanin content per feather bud area (mm2) comparing the effects of *dsFBXO32*+*hFBXO32^WT^ and dsFBXO32*+*hFBXO32^G139S^*in CNC. **F)** Diagram showing the transfected region in the trunk. **G)** Dorsal views of E13 embryo subjected to *dsFBXO32*+*hFBXO32^WT^* transfection in TNC. Melanocytes exhibit a typical differentiation in barb ridges on the transverse section (right). **H)** Dorsal views of E13 embryo subjected to *dsFBXO32*+*hFBXO32^G139S^*in TNC. On the transverse section (right), melanocytes accumulate dense black patches of eumelanin. Arrowhead highlights the expansion of vascular development into the dermis. **I)** Box and whiskers plots comparing the effects of *dsFBXO32*+*hFBXO32^WT^ and dsFBXO32*+*hFBXO32^G139S^* in TNC on feather length. **J)** Average melanin content per feather bud area (mm2) comparing the effects of *dsFBXO32*+*hFBXO32^WT^ and dsFBXO32*+*hFBXO32^G139S^*in TNC. **K)** Western blot analysis of *FBXO32* expression after 40h post-electroporation of TNC with *dsFBXO32*+*hFBXO32^WT^*or *dsFBXO32*+*hFBXO32^G139S^*. Quantification of protein intensity relative to b-tubulin. Differential protein contents comparison sums 5 embryos subjected to functional analyses in TNC and CNC, 5 embryos subjected to *dsFBXO32,* 4 embryos subjected to *dsFBXO32*+*hFBXO32^WT^,* and 4 embryos subjected to *dsFBXO32+hFBXO32^G139S^*. Bars indicate the mean; error bars, SD; one-way ANOVA with Dunnett’s multiple comparison tests (\*\**P*<0.05, \*\*\*\**P*<0.0001, ns: not significant). Scale bars: 50µm.

Second, we explored *FBXO32’s* implication in TNC-derived melanocyte differentiation. In the trunk, the NCCs fated to form melanocytes, migrate between the ectoderm and the somite along a dorsal pathway ^35^. Unilateral electroporation performed in TNC at 22ss, between somites 16 and 22, was monitored with mCherry activity to visualize the transfected ipsilateral side (Figures 1G, 1I, 1IJ, and 1K). Immunostaining with anti-FBXO32 Ab of transfected embryos 24h post-electroporation showed that mCherry-positive TNC cells accumulated FBXO32 (Figure 1L). In transfected embryos at E13, on both untransfected and transfected sides, skin appendages exhibited melanocyte differentiation, characterized by longitudinal foxy stripes in feathers and pheomelanin accumulation (Figures 1M and 1R), similar to the cranial phenotype. When subjected to *FBXO32* silencing, TNC cells gave rise to unpolarized melanocytes accumulating eumelanin (Figure 1S), resulting in hypoplastic feathers with eumelanin pigment (Figures 1T and 1U). Moreover, melanocytes with eumelanin localized adjacent to the feather soma, as we observed in *dsFBXO32*-treated CNC. We also observed an increased blood vessel density in the dermis after *FBXO32* silencing in TNC. (Figures S2C and S2D). These observations indicate that *FBXO32* controls feather development and melanocyte differentiation and regulates vessel development in the dermis at the TNC.

### *FBXO32^G139S^* in embryonic NCCs leads to melanocyte differentiation impairment

Next, we challenged to rescue the consequences of silencing the chicken *FBXO32* construct driving the cDNA expression of the wild-type form of human *FBXO32* (*hFBXO32^WT^*) to test if the human gene could normalize pigment cell development in the CNC and TNC. Twelve days after electroporation with *dsFBXO32+hFBXO32^WT^*in CNC, we observed restoration in the feather development and melanocyte differentiation (Figures 2A and 2B). We noticed a mild feather hyperplasia attributable to the strong CMV promoter driving *hFBXO32^WT^*gene expression in the developing pigment cells ^36^. Strikingly, CNC-derived melanocytes subjected to *hFBXO32^WT^* expression could restore pheomelanin production and accumulation. These results indicate that the human wild-type gene can interspecifically restore defective melanocyte differentiation and fully mitigates the adverse effects of the chicken gene silencing on feather morphology and pigment phenotype.

In parallel, we used a similar strategy to unmask the specific effect of the variant identified in the affected children, p.G139S, referred to as *hFBXO32^G139S^.* The *hFBXO32^G139S^* construct was unilaterally co-electroporated with *dsFBXO32* to interspecifically compensate for the silencing of the endogenous gene and reveal the specific impact of the mutation on the CNC and TNC differentiation. In the CNC, electroporation with the *dsFBXO32+hFBXO32^G139S^* resulted in a considerable increase in eumelanin synthesis (Figure 2C). Melanocytes exhibited disrupted polarity with their filopodia oriented towards the dermal pulp of the feather. Consistently, co-electroporated *dsFBXO32+hFBXO32^G139S^*-embryos exhibited a massive eumelanin synthesis, comparable to *dsFBXO32-*treated embryos (Figures 2D and 2E).

Furthermore, we evidenced eumelanin-pigmented cell progression towards the dermis: these cells were strongly positive for immunostaining using anti-melanoma cell-cell adhesion molecule (MelCAM) Ab. In cranial skin, immunolabelling revealed a faint MelCAM Ab immunoreactivity of epidermal *dsFBXO32+hFBXO3^WT^*-CNC cells (Figure S3A). By contrast, embryos transfected with *dsFBXO32+hFBXO32^G139S^*showed aberrant MelCAM immunolabelling in both epidermis (Figure S3Bi) and dermis (Figure S3Bii). In addition, the *dsFBXO32+hFBXO32^G139S^* melanocytes were massively observed near nerve sheaths and blood vessels, indicating an abnormal tropism towards Schwann and peri-vascular cells (Figure S3Biii and S3Biv). Our results showed that *FBXO32^G139S^*triggered aberrant MelCAM immunoreactivity and ectopic migration at the dermis, thus illustrating a possible role of the variant in melanocyte transformation and pigment shift during NC development.

In the TNC, co-electroporation of *dsFBXO32*+*hFBXO32^WT^*at 22ss restored feather polarity, pheomelanin pigmentation, and normalized melanocyte distribution in the barb ridges (Figures 2F and 2G). Conversely, the co-electroporation of *dsFBXO32+hFBXO32^G139S^* perturbed feather polarity where melanocytes formed dense aggregates with eumelanin synthesis (Figure 2H). The cells were also misoriented, projecting their extensions towards the dermis. Compared to *dsFBXO32+hFBXO32^WT^*, we showed that *FBXO32^G139S^* provoked hypoplastic feather buds with enriched melanin content (up to 20%; Figures 2I and 2J). At the protein level, on truncal skins, 40h post-electroporation, *hFBXO32^G139S^*sharply reduced FBXO32 protein accumulation, while *hFBXO32^WT^* augmented the FBXO32 expression on Western blot analysis (Figure 2K). These *in vivo* functional analyses highlight the critical functional role of *FBXO32* in CNC and TNC and the detrimental effect of *FBXO32^G139S^* on melanocyte differentiation.

### Identification of *hFBXO32^G139S^* transcriptional targets in CNC and TNC

Given the striking phenotypes yielded by the functional analysis in cranial and trunk samples, we performed gene expression profiling by RNAseq analysis on cranial and truncal skin harvested at E13. Hierarchical clustering showed two distinct features (Figure 3A) and a good correlation of gene expression between the three biological replicates. We noticed significant differences in gene expression between the conditions tested in cranial and trunk skins. Third, gene expression clusters exhibited opposite changes between cranial and truncal skins (Figure 3A).

**Figure 3:**
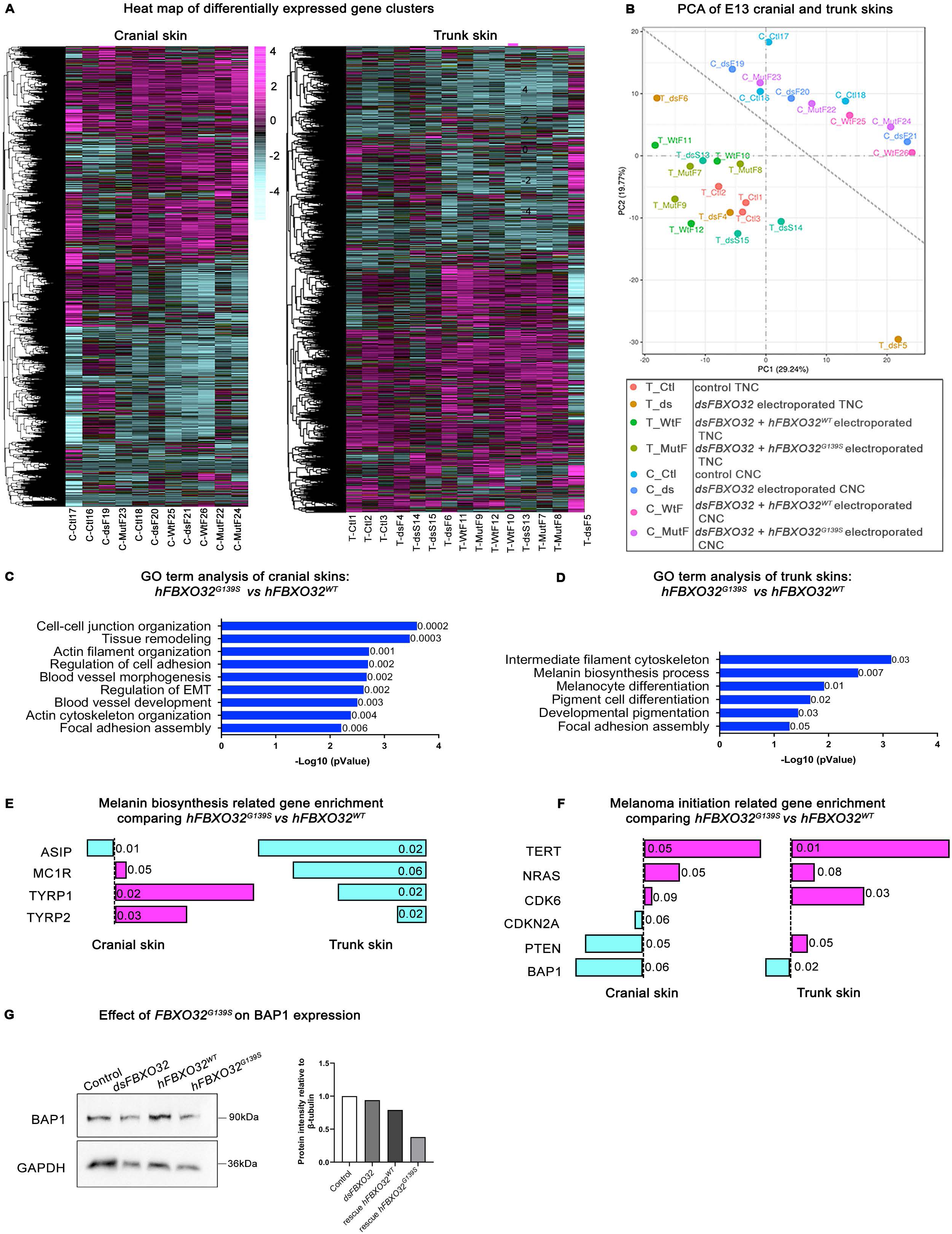
Trunk and cranial samples colonized by TNC and CNC exhibit distinct transcriptomic profiles in response to physiological or altered *FBXO32* functions. **A)** Heat map showing the hierarchical clustering of genes in the trunk and cranial E13 skin samples. **B)** PCA plot based on the trunk and cranial skin samples at E13: the two clusters indicate that cranial and trunk samples have differential gene expression patterns. **C-D)** Significantly enriched signaling pathways in cranial and trunk samples, respectively, as predicted by GO analysis. **E)** Bar chart showing differentially expressed genes related to childhood melanoma drivers. Left: Bar chart of up- and down-regulated genes related to melanogenesis in samples electroporated with *dsFBXO32*+*hFBXO32^G139S^*compared to samples electroporated with *dsFBXO32*+*hFBXO32^WT^*samples. **F)** Bar chart showing up-and down-regulated genes related to melanoma initiation in trunk samples taken from *dsFBXO32*+*hFBXO32^G139S^*-electroporated embryos compared to *dsFBXO32*+*hFBXO32^WT^-*electroporated embryos. **G)** Effect of *FBXO32^G139S^* on BAP1 expression 11 days post-electroporation by Western blot.

The principal component analysis (PCA) revealed the relative distances between all triplicates in cranial and truncal samples (Figure 3B). Cranial samples cluster appeared spatially distinct from the truncal ones. To identify the transcriptional changes due to the *FBXO32* variant, we compared genes statistically enriched or reduced in cranial and trunk samples (Figures S4A and S4B). In cranial samples, the comparison between *dsFBXO32+hFBXO32^G139S^* and *dsFBXO32+hFBXO32^WT^*, revealed 158 upregulated genes and 69 downregulated genes (Figure S4A). Signaling pathways enrichment analysis related to NC development pointed to the upregulation of *BMPR2A, EGFR,* and *BMPR1A* genes and the downregulation of *WNT4, FGF12, SHH, PTCH2, SANI1, TWIST1* genes (Figure 4SC). In parallel, the volcano plot of differentially expressed genes in trunk samples between *dsFBXO32+hFBXO32^G139S^* and *dsFBXO32+hFBXO32^WT^*showed 82 genes upregulated (among *WNT4, SOX9, FZD6, WNT7A*) and 52 genes downregulated (Figure 4SB), among which *SOX10* was where the most striking target (Figure S4D).

**Figure 4:**
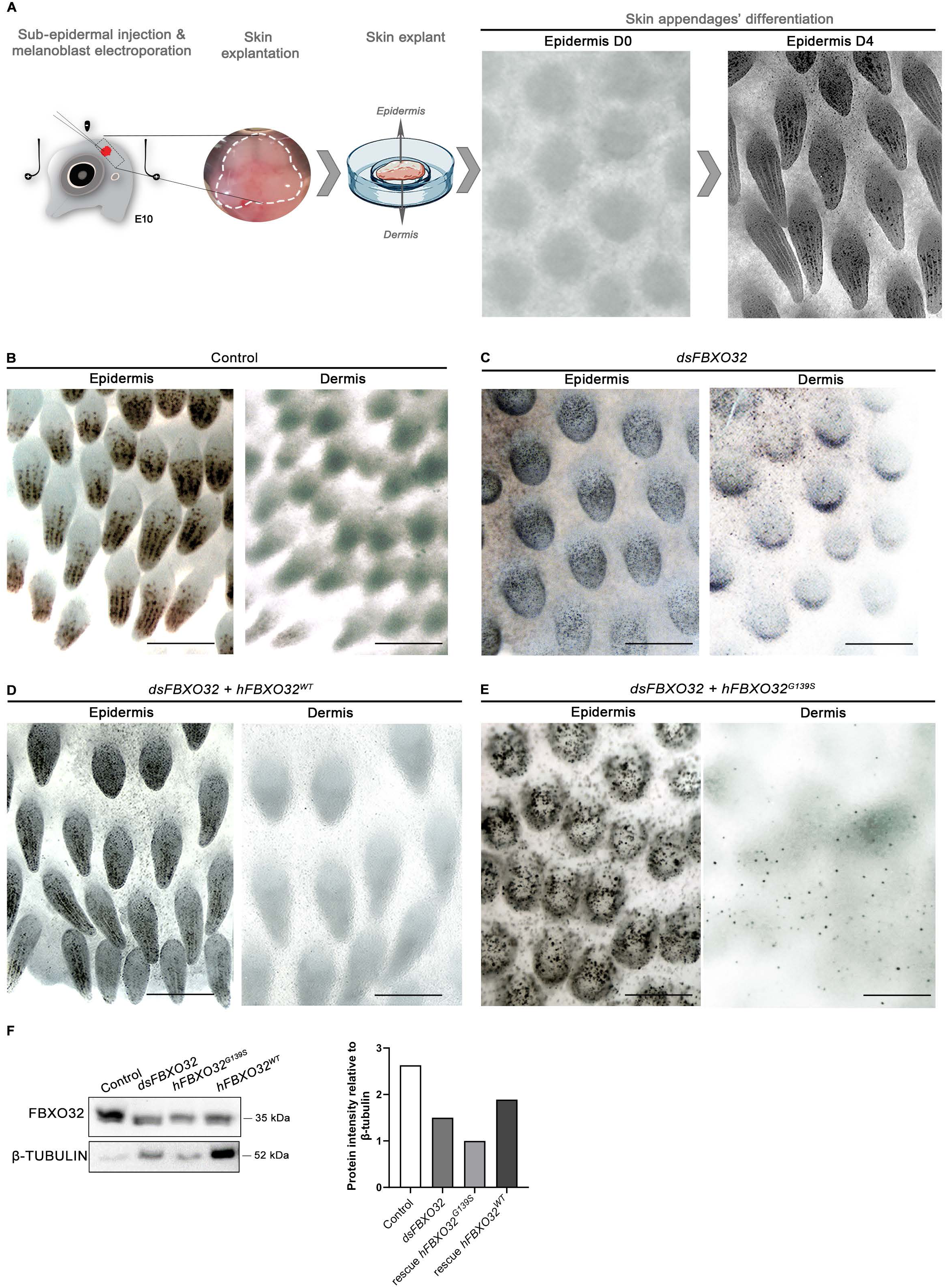
*FBXO32^G139S^* provokes melanocyte proliferation and migration toward the dermis. **A)** Diagram showing melanoblast electroporation and *ex vivo* skin explantation at E10. Exogenous nucleic sequences are micro-injected in the dermis (red). Following bilateral electroporation (considered as D0), skin explants are harvested and laid on a Petri dish, the dermal side in contact with the substratum, and maintained for four days *in vitro*. **B)** From D0 to D4, melanogenesis occurs along the medial longitudinal axis and progresses from mesial to lateral:melanocytes differentiate in the skin epidermis at D4. **C)** Skin explants subjected to *dsFBXO32* exhibited atypical melanocyte migration toward the dermis. Right, epidermis side; left, dermis side. **D)** Skin explants subjected to *dsFBXO32+hFBXO32^WT^*recover feather bud morphology and melanocyte differentiation in the dermis. **E)** Skin explants subjected to *dsFBXO32+hFBXO32^G139S^*exhibit hypoplastic feather buds and ectopic melanocyte migration toward the dermis. **F)** Western blot analysis of *FBXO32* expression in control, *dsFBXO32-, dsFBXO32+hFBXO32^WT^-, and dsFBXO32+hFBXO32^G139S^-*transfected skin explants, at D4. Quantification of protein intensity relative to b-tubulin. D: days *ex vivo*. Scale bars: 500µm.

To gain insight into the impact of the variant on biological processes, we performed a Gene Ontology (GO) enrichment analysis. We found gene enrichment associated with actin cytoskeleton organization, actin filament organization, regulation of cell adhesion, focal adhesion assembly, EMT regulation and blood vessel morphogenesis, and blood vessel development in cranial samples (Figure 3C). Similarly, in truncal samples, the mutation targeted biological processes related to intermediate filament cytoskeleton and focal adhesion assembly. Additionally, we found differential enrichment in the melanin biosynthesis process, melanocyte differentiation, pigment cell differentiation, and developmental pigmentation (Figure 3D). We previously showed that TNC *dsFBXO32-*transfected and *hFBXO32^G139S^*-cotransfected exhibited a shift in melanin synthesis. GO analysis further confirmed the significant enrichment in genes related to melanogenesis. Next, we identified the statistically enriched melanin-related genes between *hFBXO32^G139S^ and hFBXO32^WT^*. This analysis pointed to the downregulation of the *ASIP* gene in both cranial and trunk samples (Figure 3E). On its own, *ASIP* downregulation could account for the shift in pheomelanin to eumelanin synthesis. We then focused our analysis on eumelanin-related genes. In cranial samples, we found opposite changes between upregulated genes, including striking enrichment in *TYRP1, MLANA,* and *TYR*, and downregulated genes, including *DOPA* and *TYRP2,* (Figure S4E). In trunk samples, the upregulation of melanogenesis-related genes included *TYR, DOPA,* and *MCAM*, and the downregulation of *OCA2, MLANA, TYRP1,* and *TYRP2* (Figure S4F). These data illustrate how *FBXO32* could affect gene enrichment during NC development, cell migration, and melanogenesis.

Although the genomic landscape of adult melanoma is well documented, pediatric melanoma’s genomic features remain largely unknown. We evaluated gene enrichment triggered by *hFBXO32^G139S^* into adult melanoma drivers. We noticed changes in *NRAS, CDK6*, *CDKN2A*, *PTEN*, *TERT*, and *BAP1* expression in both cranial and trunk samples. Moreover, the *CDKN2A* gene, which is rare in early-onset melanoma, was downregulated in cranial samples. Interestingly, both cranial and truncal samples exhibited downregulation of the childhood melanoma driver *BAP1* (Figure 3F). This result suggests that *FBXO32* might counteract *BAP1* activity in the protein degradation interplay. In addition, Western blot analysis showed that the silencing of *FBXO32* and co-electroporation with *dsFBXO32+hFBXO32^G139S^* decreased BAP1 protein level in trunk samples 11 days post-electroporation. Together, gene expression profiling indicates that *FBXO32^G139S^* affects melanoma driver genes transcription.

Moreover, *FBXO32^G139S^* affected signaling pathways involved in cell biological processes. In cranial samples, we found upregulation in genes related to programmed cell death, including *PDCD11, PDCD6IP,* and *PIM1*, proto-oncogenes such as *FYN* and *SRC,* autophagy (*AMBRA1*), downregulation in apoptosis (*DAD1, CASP7*), cell differentiation (*TPD52*) and cell cycle (*RPRM*) (Figure S4G). Similarly, in trunk samples, we found upregulated genes associated with cell growth and proliferation (*DDIT4*), RAS oncogene family (*RAB7B*), cell death (*TP53I3*), and downregulated genes related to cell motility (*C1QTNF8*), cell metabolism (*MIGA2*) and cellular proliferation (*Ki67*) (Figure S4H). These results strengthen our *in vivo* experiments where *FBXO32^G139S^* impaired NCCs development.

The increased blood vessel density observed in *dsFBXO32+hFBXO32^G139S^*samples indicated that the variant could promote angiogenesis in the dermis where the electroporated NCCs migrated. The genes associated with angiogenesis pointed to the enrichment of *AMOT, AQP3, AGTRAP,* and *FGF12* as targets of *hFBXO32^G139S^* (Figure S2E and S2F). Additionally, we noticed enrichment inflammatory-related genes (*IL6, TRAF1, SLC7A2*), angiopoietins members of the vascular endothelial growth factor family (*ANGPTL2, AGTRAP, CYR61, VEGFA*), and cancer progression genes (*UPF1, PLAA, TRAF1, TFPI2*) were significantly enriched. Moreover, gene enrichment unique to pericyte recruitment (*PDGFβ*) and pericyte regulation (*CD44*) indicated a possible pericyte recruitment and vascular pattern formation. Even though gene enrichment was low regarding p-values, the transcriptomic profiling evidenced some angiogenesis-related and pericyte-related genes as specific targets of *FBXO32^G139S^*, which prompted the angiotropism of transfected NCCs leading to local vascular expansion.

### *FBXO32^G139S^* in melanoblasts triggers aberrant melanocyte migration in the dermis

To further characterize the effect of *FBXO32^G139S^* in melanoblasts, we set an original *ex vivo* skin explant model. In this system, we preserved the 3D tissue architecture between the epidermis and the dermis to track melanocyte differentiation in the epidermis and, at once, monitor the emergence of melanocyte aggregates into the dermis (Figure 4A). For this purpose, nucleic acids’ (RNAi and plasmids) were intradermally injected and electroporated before melanocyte differentiation at E10 (Figure 4A). Then, the skin was micro-dissected and transferred onto a non-embryo toxic Petri dish coated with serum (considered as day 0, D0). At D0, the dermis harbored unpigmented melanoblasts; the skin surface was flat and devoid of appendages (Figure 4A). From D0 onwards, feather buds started to develop and grow *ex vivo* following an anteroposterior and mediolateral pattern similar to *in vivo* development. After four days *ex vivo*, skin appendage differentiation recapitulated *in vivo* development at E13, with conspicuous melanocyte differentiation in the epidermis (Figure 4B). In this context, *FBXO32* silencing hampered feather bud growth and polarity: it altered bud diamond-shape patterns in the epidermis and triggered melanocyte migration towards the dermis (Figure 4C). The skin explants transfected with *dsFBXO32+hFBXO32^WT^* recovered feather morphogenesis and melanocyte differentiation (Figure 4D).

By contrast, when transfected with *dsFBXO32*+*FBXO32^G139S^*, skin explants exhibited severe alterations: hypoplastic feather buds with aberrant differentiation of melanocytes in the epidermis and in the dermis where they formed dense aggregates (Figures 4E). In parallel, we performed immunolabelling using anti-MelCAM Ab and DAPI staining on 3D skin explants. In the skin explant co-electroporated with *dsFBXO32+hFBXO32^WT^*, small MelCAM-positive foci represented melanocyte filopodia extending toward the skin surface in the epidermis. In contrast, in *dsFBXO32*+*FBXO32^G139S^*explants, MelCAM Ab highlighted tissue disorganization resulting from melanocyte aggregates in the epidermis and dermis layers (Figures S5A and S5B). This result confirmed the migration of melanocytes to the dermis when subjected to *FBXO32^G139S^* and was consistent with the invasive radial growth phase observed in human melanomas ^37^. Furthermore, we analyzed FBXO32 protein accumulation in the skin explants subjected to *dsFBXO32* and *dsFBXO32+hFBXO32^G139S^*. We observed a low protein detection in *FBXO32* silencing, decreased by at least half in *FBXO32^G139S^*, while *dsFBXO32+hFBXO32^WT^* correlated to higher levels of FBXO32 expression in skin explants (Figure 4F).

### *FBXO32^G139S^* regulates cell filopodia extension, lamellipodia formation, and cell aggregation in melanocytes

Transcriptomic analysis revealed gene enrichment associated with melanocyte transformation, cytoskeleton remodeling, and focal adhesion. We decided to document the impact of *FBXO32^G139S^ on* melanocyte cytoskeleton *in* vitro. After dissociation from skin explants, melanocytes were grown on fibronectin-coated chamber slides (Figure 5A) until confluence. Melanocytes isolated from control explants exhibited fibroblastic-like morphology, with bipolar polarity and distant cell connections forming a loose network on the substratum (Figure 5B, first column). Upon *FBXO32* silencing, melanocytes had reduced filopodia length, short lamellipodia protrusions, and formed cell aggregates (Figure 5B, second column). When electroporated with *dsFBXO32+hFBXO32^WT^*, melanocytes recovered their cell morphology and polarity and restored distant cell contacts (Figure 5B, third column). By contrast, when subjected to *dsFBXO32+hFBXO32^G139S^,* melanocytes exhibited altered morphology, with short-distance contacts leading to densely packed aggregates. (Figure 5B, fourth column). These results strengthened our previous observations on *in vivo* and *ex vivo* models, where melanocytes tended to be more compact when transfected with *FBXO32^G139S^*. We quantified the filopodia length in *dsFBXO32* and *dsFBXO32*+*hFBXO32^G139S^*: both conditions decreased two-fold filopodia length compared to filopodia length average in control cells (84.62µm; Figure 5C). Together, these results reflect a dynamic and detrimental cell morphology remodeling in *FBXO32^G139S^*transfected cells.

**Figure 5:**
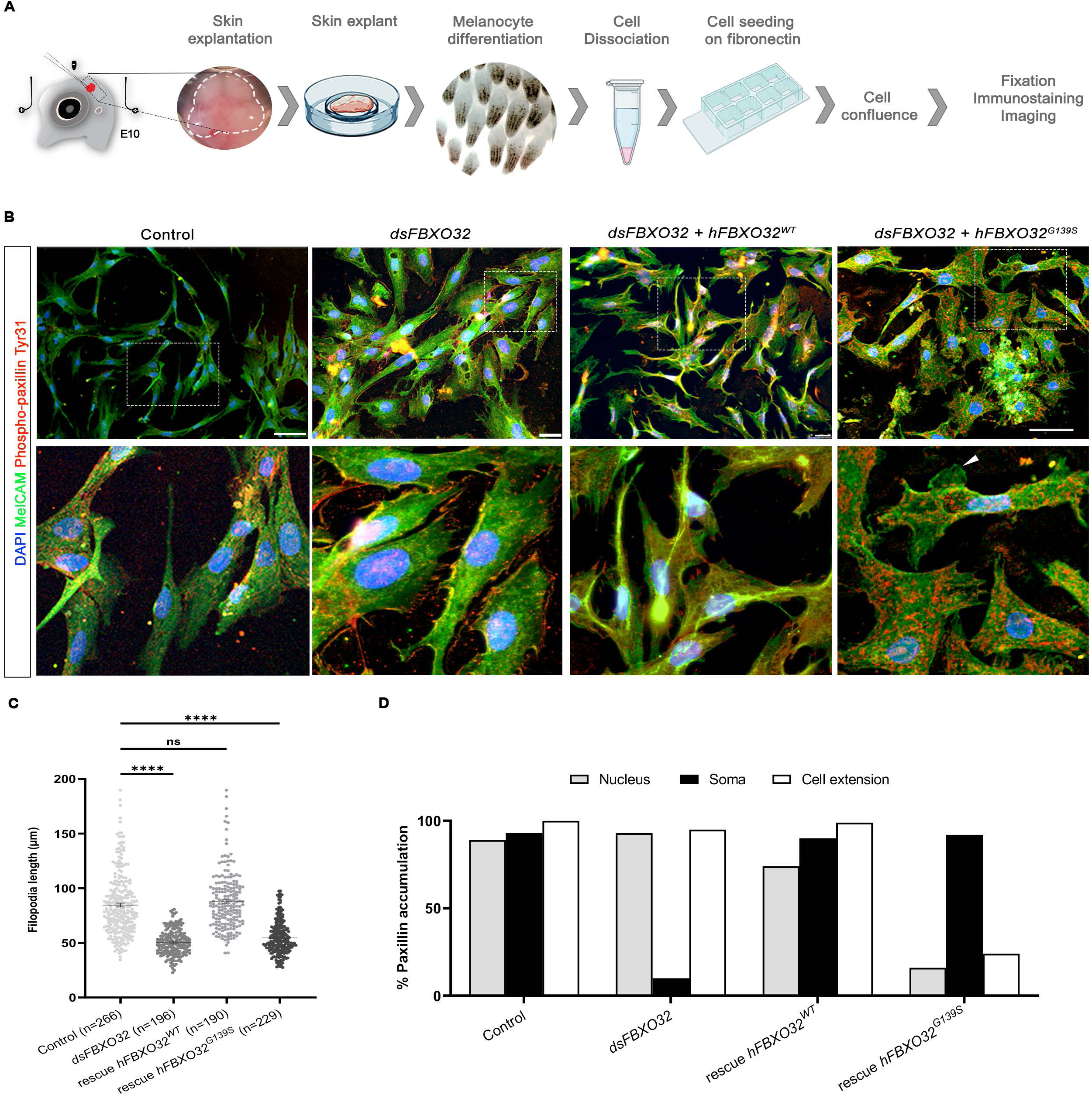
*FBXO32^G139S^* induces melanocyte transformation and cytoskeleton remodeling. **A)** Diagram showing melanocyte isolation from skin explants for *in vitro* cell culture. **B)** Confocal images of melanocytes isolated from control, *dsFBXO32-, dsFBXO32+hFBXO32^WT^-* and *dsFBXO32+hFBXO32^G139S^-*treated skins, showing their respective immunoreactivity for anti-MelCAM mAb (green) and anti-phospho-paxillin Tyr31 pAb (red). Boxes areas show high magnifications of each experimental condition showing phospho-paxillin Tyr31 accumulation. **C)** Box and whiskers plots showing quantification of filopodia length (mm) per melanocyte cells (median with interquartile range). n= number of filopodia analyzed from three independent experiments. **D)** Bar chart showing the percentage of paxillin accumulation in the nucleus, soma, and cell extensions for all experimental series. Bars indicate the mean; error bars, SD; one-way ANOVA with Dunnett’s multiple comparison tests (\*\**P*<0.05, \*\*\*\**P*<0.0001, ns: not significant). Scale bar: 50µm.

### *FBXO32* gene inactivation and *FBXO32^G139S^* increased paxillin phosphorylation recruitment

Our transcriptomic analysis suggested gene enrichment in focal adhesion assembly (Figures 3C and 3D). Focal adhesion assembly was assessed by immunostaining using an anti-phospho-paxillin Tyr31 Ab (Figure 5B). Paxillin is a crucial focal adhesion component, which functions as a scaffolding protein for assembling the multi-protein complex and facilitating intracellular signaling ^38,39^. Moreover, paxillin contributes to cell migration by coordinating cell assembly and disassembly at the front and rear by regulating focal adhesion dynamics ^40^. In control cells, we found phospho-paxillin accumulation in the nucleus, cell soma, and cell extensions (figure 5D, first column, and 5D). Cells subjected to *FBXO32* silencing showed a high accumulation of phospho-paxillin Tyr31 in cell extensions and translocation to the nucleus (Figure 5B, second column, and 5D). When subjected to *dsFBXO32+FBXO32^WT^*, cells recovered phospho-paxillin Tyr31 accumulation in the cell soma, the nucleus, and extensions (Figure 5B, third column and 5D). By contrast, in *dsFBXO32*+*FBXO32^G139S^*transfected cells, phospho-paxillin mainly accumulated in the cell soma (Figure 5D, fourth column). From these findings, we infer that *FBXO32^G139S^* targets phospho-paxillin and compromises melanocyte focal adhesion assembly, resulting in aberrant cell aggregation and disoriented navigation.

### *FBXO32* silencing and *FBXO32^G139S^* provoke actin disassembly

To further characterized *FBXO32^G139S^*’s implication in actin filament assembly, we examined actin filament organization by phalloidin staining. In normal melanocytes, actin filaments form polarized stress fibers underlying cell morphology and scaffolding their extensions (Figure 6A, first column). By contrast, in *dsFBXO32*-transfected melanocytes, the actin filament architecture was broken, compromising stress fibers and cell morphology (Figure 6A, second column). In melanocytes subjected to *dsFBXO32+hFBXO32^WT^,* cells recovered actin cytoskeletal structure and maintained actin polymerization (Figure 6A, third column). However, when subjected to *dsFBXO32*+*FBXO32^G139S^*, melanocytes exhibited actin filament remodeling (Figure 6A, fourth column). Overall, quantitative analysis showed the disruption of actin filament length size in *dsFBXO32-* and *dsFBXO32+hFBXO32^G139S^*-treated melanocytes (Figure 4B). The *dsFBXO32+hFBXO32^G139S^*-treated melanocytes exhibited a significant reduction in phalloidin accumulation compared to the control (Figure 6C). The transcriptomic analysis further documented the role of *FBXO32^G139S^*on the cytoskeleton remodeling in skin samples by showing specific gene enrichment in tight junctions (*TJP2, TJP3*), laminin (*LAMA3*), integrins (*ITGA8, ITGA1, ITFG1*), microtubule (*BCAS3*), membrane proteins (*SMIM5, BCAS3*), extracellular matrix (*TNC*) and fibronectin (*FNDC4*). We also noticed the downregulation of genes involving actin binding *KPTN*, tubulin *TUBB2A*, collagen *COL27A1*, cellular matrix *SMARC5*, and transmembrane protein *TMEM116* in cranial and trunk skin samples (Figures 6D and 6E). These results reveal that *FBXO32^G139S^* affects cell morphology at both single-cell and cell population levels by impairing cytoskeleton dynamics and intercellular interactions.

**Figure 6:**
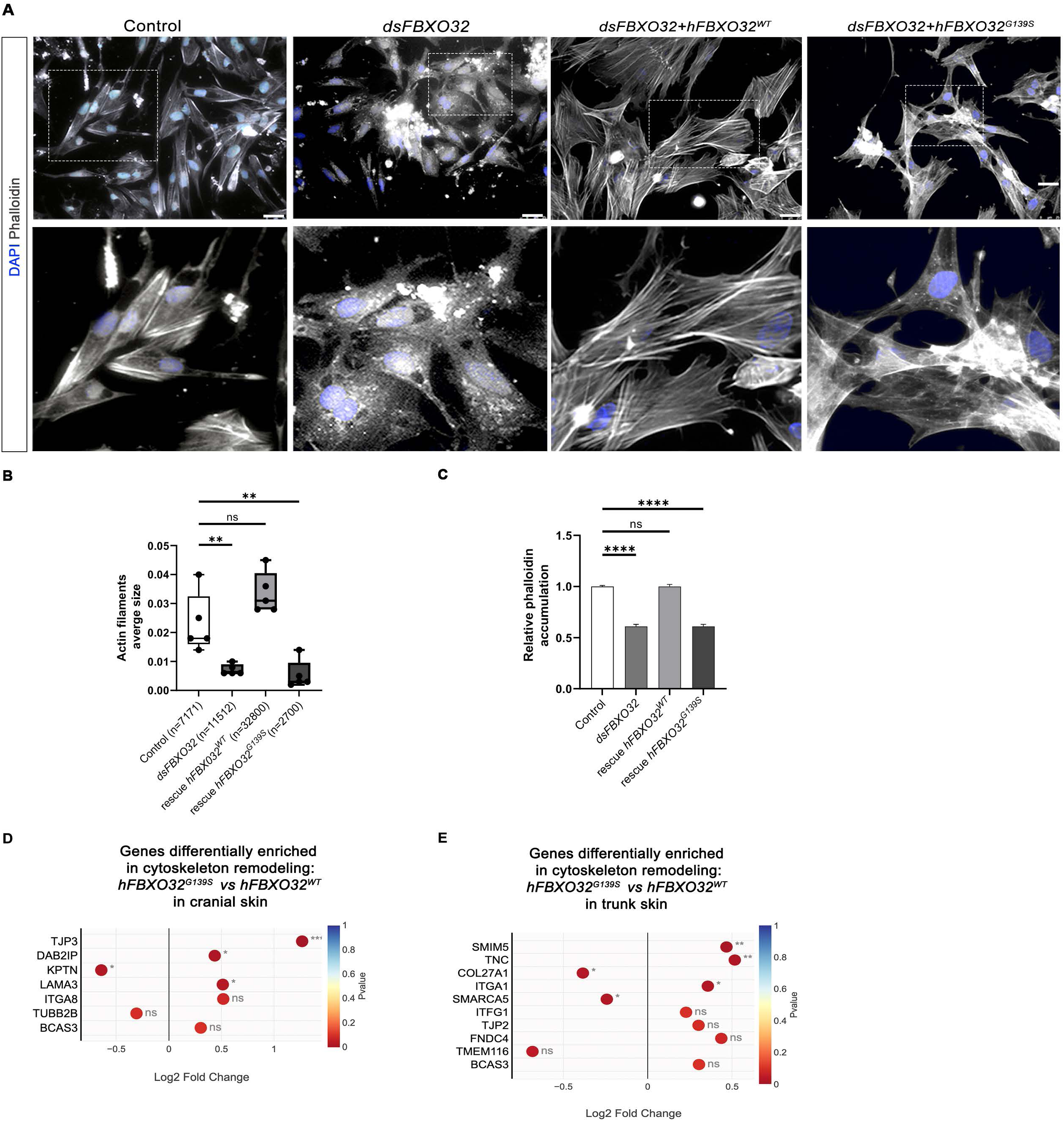
*FBXO32^G139S^* induces actin filaments disassembly. **A.** Confocal images of melanocytes isolated from control, *dsFBXO32-, dsFBXO32+hFBXO32^WT^-* and *dsFBXO32+hFBXO32^G139S^-*treated skins, showing actin filament morphology following Phalloidin staining. Boxes areas show high magnifications of each experimental condition. **B)** Box and whiskers plots showing quantification of actin filaments per field and normalized on control series. Each dot represents the average of five independent experiments (n= total number of cells analyzed per experiment). **C)** Bar chart showing relative accumulation of phalloidin in different conditions. **D-E)** Scatters plots summarizing the most differentially expressed genes in cytoskeleton remodeling between *dsFBXO32+hFBXO32^WT^-* and *dsFBXO32+hFBXO32^G139S^-*treated cranial and trunk skins, respectively. The circle color indicates the significant level with the p-value <0.05. Bars indicate the mean; error bars, SD; one-way ANOVA with Dunnett’s multiple comparison tests (\*\**P*<0.05, \*\*\*\**P*<0.0001, ns: not significant). Scale bar: 50µm.

## DISCUSSION

Clinical data from pediatric cases evidence primary melanoma cases from the head, neck, and lower extremities ^6^. Here, we analyzed melanocyte migration and differentiation at two anatomical levels, the CNC and TNC. For this purpose, we set developmental models to unmask the impact of a childhood melanoma candidate gene *FBXO32,* carrying a *de novo* mutation *FBXO32^G139S^* on melanocyte development. First, we validated *FBXO32* expression pattern in NCCs, from the onset of their migration, and later, during their differentiation into melanocytes, in the cephalic and trunk regions. Then, the RNAi approach documented how crucial *FBXO32* is for developing melanocyte lineage. To investigate the molecular mechanism by which *FBXO32* in the CNC and TNC affects the developmental program of melanocyte differentiation, we silenced its activity before NCC delamination and in the NC-derived melanocytes using RNAi-mediated gene silencing ^41^. The dsRNA driving silencing inhibited *FBXO32* activity in pre-migratory CNC with long-term effects at E13. We showed that *FBXO32* regulates melanocyte differentiation, feather polarity, and increased blood vessel density. We also investigated whether *FBXO32* affects melanocyte development in TNC. *FBXO32* silencing in the TNC correlates with CNC features: melanocyte proliferation, aberrant feather polarity, and blood vessel increase. Our functional studies established that *FBXO32* is critical in melanocyte differentiation during NC development.

Next, we challenged the phenotype resulting from *FBXO32* silencing by co-transfecting either *hFBXO32^WT^* or its variant *FBXO32^G139S^*. We showed that *hFBXO32^WT^* could inter-specifically alleviate and offset the defective melanocyte differentiation yielded by *dsFBXO32*. When this strategy was applied to explore the specific impact of *FBXO32^G139S^*, we unmasked the deleterious effects yielded by the mutation on melanocyte differentiation and migration, and documented its possible oncogenic effect. Rescue experiments demonstrated overexpression with *hFBXO32^WT^* restored melanocyte development in both NC regions. Strikingly, the rescue experiments carrying the human variant *FBXO32^G139S^* evidenced an aberrant melanocyte differentiation, pigmentation, and cell polarity. Additionally, melanogenesis in experimental embryos led to aberrant MelCAM accumulation in the dermis and ectopic localization in Schwann and pericyte cells. Normal melanocytes barely express this cell adhesion molecule, whereas melanoma cells exhibit an augmented MelCAM accumulation. The expression of MelCAM is known to correlate with an aggressive invasive phenotype, and its upregulation is strongly associated with melanoma progression ^42,43^. Although we observed a basal MelCAM protein accumulation in control skin explant and cell culture, the silencing of *FBXO32* and *FBXO32^G139S^* intensely enhances MelCAM accumulation in both systems. This observation strongly indicates that *FBXO32* dysfunction induces melanocyte transformation towards melanoma phenotype.

Angiogenesis experiments of coculture tumor cells in the chick chorioallantoic membrane hint at invasive melanoma cell recruitment called pericyte mimicry ^44,45^. In our experiments, both *dsFBXO32* and *FBXO32^G139S^* lead to an increased angiogenesis at the expense of sub-ectodermal tissues. Moreover, the CNC cells have the unique capacity to form both melanocytes and perivascular cells. The expansion of the vasculature highlights the angiotropism of CNC-derived melanocytes exacerbated by *FBXO32^G139S^*transfected CNC-derived melanocytes. These results suggest that such angiotropism, could strengthen the noxious impact of *FBXO32^G139S^* by favoring the systemic dissemination of transformed melanocytes. The coincident enrichment of genes implicated in angiogenesis, like *AMOT, AQP3,* and *FGF12* in both cranial and trunk skins, strengthen these observations. We also detected gene enrichment unique to pericyte recruitment (*PDGFβ*) and regulation (*CD44*). These results support the critical role of *FBXO32* in provoking pericyte mimicry in transformed melanocytes.

We seek novel transcriptional signatures associated with *FBXO32^G139S^.* We provided some evidence of the role of *FBXO32* in melanogenesis in both cranial and trunk series. Hierarchical clustering and PCA analysis identified two distinct cluster signatures between the cranial and trunk transfected samples. Besides, our findings demonstrate similar phenotypes from *FBXO32^G139S^* expression in cranial and trunk samples. Two regulatory events associated *FBXO32* with eumelanin synthesis. First, RNA-seq data analysis of melanin biosynthesis-related genes showed that *FBXO32^G139S^* upregulated the genes *TYRP1* in cranial skin and *Tyr* in trunk skin samples. Second, *FBXO32^G139S^* downregulated *ASIP* in both cranial and trunk compartments. The genes *MC1R, ASIP,* and *TYRP1* are significant pigmentation regulators of eumelanin biosynthesis. The *ASIP* protein, an *MC1R* antagonist, inhibits eumelanin synthesis by directly competing with α-MSH for binding to MC1R, resulting in pheomelanin synthesis ^46^. *FBXO32^G139S^* may act upstream of *ASIP*, indirectly causing *ASIP* downregulation and *TYRP1* upregulation, stimulating eumelanin synthesis (Figure S6). This study establishes *FBXO32^G139S^* role in melanogenesis during NC development.

We investigated the role of *FBXO32* in the early onset of melanoma by analyzing the epistatic relationship between childhood melanoma candidate genes. Our RNAseq data indicates that *CDKN2A* is detectable during the development of normal melanocytes. Still, transcriptomic analyses provide no evidence of differential *CDKN2A* enrichment expression in the RNA-seq data between TNC series. This finding supports the notion that *CDKN2A* is rare in childhood melanoma ^6–9^. Second, we established *RNAS* and *CDK6 gene* upregulation in the expression of *hFBXO32^WT^*and *hFBXO32^G139S^*, respectively. Third, our model reveals that *hFBXO32^G139S^* upregulated the telomere reverse transcriptase (*TERT)* gene in telomere maintenance. The upregulation in the TERT gene could explain melanocyte proliferation and may predict clinical outcomes of ongoing melanoma processes ^47^. Our study also identified that *hFBXO32^G139S^* downregulated the gene *BAP1* in cranial and trunk skin samples. *BAP1* is a deubiquitinate protein with roles in numerous cell processes, including DNA damage response, cell cycle regulation, cell growth, metabolism, and the regulation of inflammatory responses ^48–51^. *BAP1* loss-of-function germline mutation is a tumor predisposition gene in childhood melanoma ^19,20^. Immunohistochemical studies showed decreased nuclear *BAP1* localization with a dense, possibly pigmented, clonal sheet-like area within *BAP1* inactivated nevus ^51^. Our RNA-seq data indicates that *hFBXO32* downregulates the *BAP1* gene and regulates other genes involved in the ubiquitin protein degradation system (Figure S6).

*In vivo* phenotype analysis revealed a possible melanocyte transformation in delaminating NCCs. We uncovered that *FBXO32^G139S^* favors the proliferation and migration of melanocytes to the dermis recapitulating a possible invasion process. Recently a study of *FBXO32* in adult melanoma described a decreased migration of melanoma patients’ cells and 501Mel cell line in Boyden chambers assays due to the inhibition of *FBXO32* ^26^. Moreover, forcing *FBXO32* expression using a lentiviral vector in cells expressing mutant *BRAF^V600E^* on a wild-type *NRAS* stimulated SKmel28 melanoma migration ^26^.

Here, we analyzed the respective roles of *FBXO32* and the *FBXO32^G139S^*variant during NCC and melanocyte development without activating other melanoma drivers. In addition, we set a 3D-ex-vivo system that preserved the critical dermis-epidermis interactions, and allowed melanogenesis and skin appendage morphogenesis to be mimicked and recapitulated in a dish: the same samples were followed longitudinally over time. This system allowed us to track the fate of melanocyte lineage at both the epidermis and dermis levels and to follow the back-migration to the dermis of *FBXO32^G139S^*-transfected melanocytes, preceding their metastatic dissemination. Our study indicates that on its own, *FBXO32^G139S^* can trigger a possible melanocyte transformation during the development of melanocyte lineages.

Cell migration is a complex process applying adhesion, protrusion, and contractility. It relies on cell polarization and projection of cell extensions along with bonding at the leading edge and coordinated with the disassembly at the cell’s rear end ^52^. Paxillin is a scaffold/adapter protein contributing to cell migration by regulating focal adhesion dynamics ^38,53,54^. Previous work reported that paxillin phosphorylation is essential for the assembly of focal adhesion and lamellipodia protrusions formation ^55,56^. Our study used a mAb against phospho-paxillin (Tyr31) for immunolabelling transfected cells with the *dsFBXO32* condition. We observed an unexpectedly high accumulation of phospho-paxillin Tyr31 in lamellipodia protrusions and cell soma in transfected cells with *dsFBXO32* and *hFBXO32^G139S^*, respectively. We suggest phospho-paxillin Tyr31 could be *FBXO32* substrate, thus explaining phospho-paxillin accumulation when *FBXO32* is silenced or mutated. In phospho-paxillin mutants, the increased lamellipodia protrusions and destabilized actin organization accounted for disrupted focal adhesions ^55^. Here, we uncovered actin filament disassembly in *dsFBXO32* and *FBXO32^G139S^*-transfected cells, which coincided with significant gene enrichment in cytoskeleton architecture genes. This result could explain the abnormal migration of differentiated melanocytes observed *in vivo* and *ex vivo* and their propensity to aggregate. Furthermore, we observed increased phospho-paxillin translocation to the nucleus in *dsFBXO32,* which suggests a possible implication of paxillin in promoting DNA synthesis and cell proliferation ^57^. Our finding raises the implication of *FBXO32* in impairing the cytoskeleton assembly and protrusion formation.

Childhood melanoma is a rare disease, and the current knowledge relies on adults’ genetic studies. Treating children do not benefit from appropriate specific guidelines. Generally, young patients follow the same principles as adults ^58^; the issue concerns limited access to clinical trials and new drugs, dramatically altering the natural history of advanced melanomas. In the melanoma field, this is the first functional analysis to date where *in vivo, ex vivo,* and *in vitro* heterospecific models are specifically designed and used to understand the biological role of *FBXO32* in NC lineages. *FBXO32* loss-of-function promotes cell proliferation and migration, suggesting the tumor suppressor potential of *FBXO32.* Even though the interplay between *ASIP* and *BAP1* would need further analysis, our transcriptomic data raise the intriguing prospect of *FBXO32* as a novel tumor suppressor gene regulating the balance between *ASIP* and *BAP1* and shed light on the role of *FBXO32^G139S^* mutation. We propose *FBXO32* as a tumor suppressor gene that may prevent pediatric melanoma’s onset, while the variant *FBXO32^G139S^* may promote melanocyte transformation during development.

## ACKNOWLEDGMENTS

This project was supported by Centre National de la Recherche Scientifique (CNRS), fellowships from Ligue Contre le Cancer (IP-SC-13603, IP-SC-15666, IP-SC-13587, and IP-SC-17131), and grants from the Foundation for Medical Research (No FRM-DEQ20170839116 CREUZET) and the National Institute for Health and Medical Research (No INSERM (19CR055-01)/CREUZET Sophie). We also thank the Imagif campus platform (I2GB, CNRS, Gif-sur-Yvette, France) for plasmid sequencing and subcloning and Emmanuel Bruet for the critical manuscript reading. The authors thank the Doctoral School of Oncology (No 582) for the constant support for the project.

## AUTHOR CONTRIBUTIONS

A.V. and C.G. performed *in vivo* silencing experiments. A.V. performed *in vivo, ex vivo* silencing, and rescue experiments. A.V. performed 2D *in vitro* cell culture. A.V. generated cranial and skin samples for RNAseq analysis. T.G. performed RNA extraction for RNAseq analysis and WB experiments. A.V. generated immunofluorescence data from skin explants samples and cell culture. A.V. and S.C. analyzed the RNAseq database. A.V. and S.C. interpreted the results and wrote the original draft for the manuscript. S.C. designed and supervised the work. All authors read, commented on, and proofed the final manuscript.

## INCLUSION AND DIVERSITY

We support inclusive, diverse, and equitable conduct of research.

## MATERIALS AND METHODS

### Animals

The chick embryo was used as an experimental model to unravel the unknown biological significance of the melanoma candidate gene. Fertilized eggs were delivered every two weeks to the laboratory. For all experiments, eggs were incubated at 38.5±0.5°C until reaching the desired developmental stage. Embryos were staged according to the number of somite-pair (somite-stage, ss) or embryonic developmental day (E). Experiments were performed at 7ss or 22ss for *in vivo* electroporation of pre-migratory CNC or TNC, respectively. To access and manipulate the developing chick embryo, a hole is made in the chamber air to pull down the vitellus and open the shell without damaging the embryo. Then, a window is made by removing a small piece of the shell, and after embryo manipulation, the window is sealed with a transparent tape before placing the egg again in the incubator at 38.5±0.5°C. Embryos were allowed to develop until reaching the desired stage of development. Cranial and skin E13 electroporated embryos at 7ss or 22ss, respectively, were used to generate the RNAseq dataset analyses. Embryos were immediately used (RNA-seq) or fixed in 4% paraformaldehyde (PFA) in PBS at 4°C overnight for phenotyping or immunofluorescence. Experiments of undifferentiated melanoblasts were performed by *ex vivo* electroporation on skin explants at E10. All animals’ procedures *in vivo, ex vivo* electroporation, melanocyte culture, and RNAseq were performed by France and EU legislation and guidance on animal use for bioscience research.

### Chicken gene sub-cloning and constructs preparation

For silencing experiments, the chicken sequence encoding for *FBXO32* was sequenced and subcloned at the Imagif campus platform (I2GB, CNRS, Gif-sur-Yvette, France). Loss-of-function experiments were carried out using the plasmid construct PCR-TOPO-*FBXO32*. Total RNAs from *Gallus gallus* tissues were extracted using the Qiacube automation robot (Qiagen) using the RNeasy Plus Mini kit (Qiagen). The corresponding cDNAs were synthesized by reverse transcription using the Maxima H Minus First strand DNA synthesis kit (Thermo Scientific). Two specific primers have been designed. The sequence of interest was amplified with the high-fidelity polymerase PFX (Invitrogen). This amplification product was then subcloned into the vector pCRII_TOPO_TA using the TOPO® TA Cloning® Kit (Invitrogen). The subcloned sequence was validated by sequencing.

For the rescue experiment, *dsFBXO32* was co-electroporated with the construct driving either the human wild-type gene (*hFBXO32^WT^*) or the mutated gene (*hFBXO32^G139S^)*. We used the pcDNA3.1 + C-eGFP plasmid carrying the human genes *hFBXO32^WT^* and *hFBXO32^G139S^*. Plasmids used in this study were controlled by a CMV promoter. The CMV promoter yields a powerful expression of the exogenous sequence over 72h. Even though the plasmid is transiently active, its activity is sufficient to durably impact the fate of the cells until their final differentiation ^59^.

### Nucleic acid preparation

Gene silencing and rescue experiments were performed by electroporation *dsRNA* and human cDNA sequences in the chick embryos. In chicks, this technique results in highly specific gene inactivation, and such an unspecific effect has never been evidenced ^41,60^. Sense and antisense strands of RNA were synthesized from the cDNA encoding for *FBXO32* targeted gene ^41^. First, plasmid DNA purification was done using QIAfilter Plasmid Midi kits (QiAGEN). We followed the manufacture protocol, which consists of (1) a bacteria-transformed pellet resuspended in lysis buffer, (2) adsorption of DNA into the QiAGEN membrane, and (3) washing and elution of the plasmid DNA. Then, 10 to 20 μg of the plasmid containing the cDNA of the interest gene was linearized by restriction enzymes for 2h30 at 37°C Linearized cDNA enabled *in vitro* transcription of antisense and sense strands. Synthesis of RNA sense and antisense strands was done by (1) invitro using the matching RNA polymerases for 2 hours at 37°C, (2) DNA template degradation by the DNaseI free (Promega), and (3) RNA strands were precipitated by LiCl (0.5M), EDTA and EtOH 100% and cooling at −20°C, overnight. The RNA was isolated and stored at −80°Cin 25μL DEPC-H2O. The stoichiometry of RNA single strands was calibrated before annealing at 95°C for 5 min. After the elimination of the cDNA template, single strands were purified and annealed for 5 min at 95°C. The *dsRNA* against *FBXO32* was used at 0.3 µg/µl to yield rapid and efficient silencing ^61^. In the control series, solutions of non-annealed sense and anti-sense RNA strands of the targeted genes were transfected at the same concentration as in the experimental series. For rescue experiments, *dsFBXO32* RNAs were co-transfected with the human wild-type and human mutated *hFBXO32^G139S^*, cDNA at the concentration of 1.25μg/μl.

### Gene silencing and rescue experiments in avian embryos

Before electroporation, nucleic acid solutions were contrasted with 0.01% Fast Green solution in PBS (Sigma) to control the injection site carefully. Electroporation was achieved at the CNC using a triple electrode system placed on the vitelline membrane with two cathodes flanking the development head with a gap of 5 mm and the anode facing the anterior neuropore ^62^. This electrode placement generates a triangular electric field yielding the bilateral dispersion of nucleic acid sequences in the target region. ^61^ At the trunk level, electroporation was carried out using a double electrode system, with the anode flanking the transfected side and the cathode flanking the contralateral untransfected side, the latter being used as an internal control. Electroporation was achieved for both TNC and CNC by delivering five pulses of 27 volts using the electroporator system (ECM 830 BTX, Harvard Bioscience). After electroporation, eggs were sealed and re-incubated at 38.5±0.5°C until the stage required for analysis was reached. To follow the fate of migratory NCCs transfection, we initially electroporated with a pCAGGS-mCherry plasmid (Addgene n°41583) in CNC or TNC cells at 7ss and 14ss, respectively.

Samples were collected at E13 for phenotyping analysis and fixed in 4% PFA. For RNAseq analysis, cranial and trunk skin samples were collected at E13 and harvested in TRIzol (Sigma Aldrich). Each analysis was performed in triplicates. E13 embryos were imaged under a stereomicroscope Leica Wild M10 equipped with a digital camera micro-publisher 3.3 RTV and digitalization using Q-capture 6.0 software.

### *Ex vivo* electroporation on skin explant

Undifferentiated melanoblasts were transfected in the cranial skin at E10. Briefly, the molecules of interest (RNAi) or cDNA human rescue sequences *hFBXO32^WT^* or *hFBXO32^G139S^* were injected into the dermis of the dorsal cranial skin, followed by electroporation using a triplex of electrodes. In this context, exogenic nucleic acids were transferred exclusively to the melanocyte precursors in the dermis, while the epidermis corresponding to keratinocytes and where melanocytes differentiate left untransfected. The skin subjected to electroporation was explanted using micro-scissors for iridectomy and transferred into a non-toxic Petri - Nunc IVF Dish (ThermoFisher). The 3D skin explant culture system supports the differentiation of skin appendages and the development of feather buds *ex vivo*. Skin appendages develop in 4 days according to anteroposterior and mediolateral body axes and exhibit a pattern of pigmentation recapitulating the *in vivo* melanogenesis. Therefore, we tracked melanocyte differentiation on Day 4. After melanocyte differentiation on a dish, skin explants were either fixed on 4% PFA for immunostaining analysis or directly used for melanocyte cell culture.

### 2D Melanocyte cell culture

Melanocyte cells were isolated from the 3D *ex vivo* transfected skin explants. After micro-dissection, the epidermis was separated from the dermis and incubated in TrypLE^TM^ recombinant enzyme (ThermoFisher) for 1h and subsequently dissociated mechanically with a pipette tip. The resulting tissue and dissociated cells were cultivated on Lab-Tek chamber slides previously coated with fibronectin (20μm/ml; F1141, Sigma-Aldrich) in 300μl of Dulbecco’s Modified Eagle Medium/Nutrient Mixture F-12 (DMEM/F12, Gibco^TM^) + 5 % goat bovine serum (GBS, Sigma Aldrich) + 1% penicillin-streptomycin (10 000 U/ml, Gibco^TM^) + 1% Amphotericin B solution (Sigma Aldrich). 2D cell cultures were maintained in cell culture incubator Midi 40 (3404, ThermoFisher) until reaching confluence at 37°C and 5% CO_2_. Cells were fixed in 4% PFA for immunostaining analysis.

### *In situ* hybridization

Whole-mount and fixed-paraffin section *in situ* hybridizations were prepared as previously described ^63^ using the following probes: *FBXO32* (Imagif campus platform, I2BC). Fixed embryos in 4% PFA were washed in PBS 1X containing 0.1% Tween-20 (PBST). Embryos were dehydrated in a graded series of 25% and 50% methanol in PBST and incubated in 100% methanol for a cold shock at − 20°C for 30 min. Then, embryos were progressively rehydrated in methanol + PBST for 10 min and rinsed twice in PBST for 10 min. Embryos were digested with 20μg/ml proteinase K (4min, 14min, 1h for 4ss, 22ss, and 26ss, respectively) and post-fixed in 4% PFA + 0.1%glutaraldehyde for 20 min each. Prehybridization was performed by incubation with a hybridization buffer for 1h at 70°C. Then, embryos were incubated with the corresponding DIG-labeled riboprobes diluted in 1 μg/mL hybridization buffer at 70°C overnight. On the second day, riboprobes were harvested and stored at −20°C, which can be reused up to 4 times. The embryos were rinsed with pre-warmed hybridization buffer for 45 minutes, twice, then in MABT (2M maleic acid, 5M NaCl, 0.1% Tween-20, pH 7.5). Non-specific antibody binding was prevented by incubation in MABT with 2% blocking reagent and 20% goat serum for 60 minutes. Then embryos were incubated overnight at RT in AP-conjugated anti-DIG antibody (Roche, 1:1000 diluted in the blocking buffer). On the third day, the embryos were rinsed in MABT for 3h before a half-hour treatment with NTM (100 mM Tris, 100 mM NaCl, 50 mM MgCl2, pH 9.5). We reopened cephalic vesicles for embryo 24ss to avoid the non-specific accumulation of the staining solution. The hybridization staining was developed in 20 μL/1mL of NBT/BCIP (Roche) at 37°C and protected from light. Whole embryos *in situ* hybridization were imaged using a Leica MZFLIII fluorescence stereo zoom microscope; digitalization was done with the Infinity Analyze 6.3.0 software.

Paraffin sections of 7μm-thick were used for *in situ* hybridization with a DIG-UTP-labelled probe in a 1μg/mL hybridization buffer. First, the sections were deparaffined by 3 times toluene baths, progressive rehydrated in graduate ethanol dilution (100%, 90%, 70%, and 30%), then digested with 20mg/mL proteinase K for 14 min, post-fixed in 4% PFA and rinsed with PBS + saline-sodium citrate (SSC) buffer (3M NaCl, 0.3 M Sodium citrate), pH 7.5. After overnight hybridization at 70°C with the corresponding riboprobe, the slides were washed twice in 50% formamide + 1X SSC at 65°C, then in MABT, pH 7.5 at room temperature (RT). Nonspecific antibody binding was blocked by incubation in MABT with 20% goat serum and 2% blocking reagent for 90 minutes. The blocking solution was replaced with an anti-DIG antibody (1:2000) diluted in MABT and 20% blocking reagent on each slide and covered with a glass slide for overnight incubation at RT. The following day, slides were rinsed in MABT for 3h before a half-hour treatment with NTM. NBT/BCIP diluted in NTM was finally applied and kept in a humid chamber at 37°C until the revelation. Slides were mounted in Aquatex (Merck). *in situ* hybridization sections were imaged using transmitted light under the EVOS M5000 microscope (Invitrogen).

### Western blot

For Western blot analysis: 40h post-electroporated TNC samples, 11 days post-electroporated TNC samples and D4 cranial skin samples were collected from control, *dsFBXO32-*, *dsFBXO32+hFBXO32^WT^-,* and *dsFBXO32+hFBXO32^G139S^*-electroporated samples. Dissected tissues were rapidly chilled into dry ice and then at −80°C.To obtain total protein extracts, T-PER^TM^ tissue protein lysis buffer (Thermo Scientific™), supplemented with protease and phosphatase inhibitor cocktail (Thermo Scientific™), was added to the samples at 4°C. Samples were then manually homogenized using plastic pistons. Lysates were centrifugated for 5 min at 10 000 rpm; supernatants were transferred into new tubes and stocked at −80°C while pellets were thrown. Determination of protein concentration was determined using the Pierce BCA Protein Assay kit (Thermo Scientific™). An equal amount of total proteins (5-10 µg/lane) was separated by SDS-PAGE, transferred onto nitrocellulose membrane (Amersham), and Western blots were then conducted using standard procedures. Primary and secondary antibodies used: FBXO32 (Abcam, Ab168372), GAPDH (Thermo Fisher, MA5-15738), and alpha-tubulin (Active Motif, 39527). Horseradish peroxidase-conjugated anti-rabbit or anti-mouse antibodies were from Invitrogen. Protein detection was performed using an enhanced chemiluminescence kit (Bio-Rad). Densitometric protein quantification was done using Fiji software (National Institutes of Health) ^64^.

### Immunolabelling

FBXO32 detection in 22ss trunk electroporated whole-embryos with mCherry reporter gene was evidenced by immunohistochemistry using anti-FBXO32 (Abcam) monoclonal antibodies (Abs) and Phalloidin staining. Incubation with fluorophore-conjugated secondary Abs AlexaFluor-488 was carried out overnight at 4°C.

FBXO32 protein accumulation on 4ss, 10ss, and 24ss untransfected whole-embryos was evidenced by immunocytochemistry using anti-FBXO32 (Ab168372, Abcam) monoclonal antibody (mAb), anti-HNK1 mAb (CD57; Sigma), and staining with Alexa Fluor™ 633-conjugated phalloidin (Invitrogen™). Briefly, whole embryos were washed and fixed at room temperature for 20 min with 4% PFA, dehydrated gradually in methanol baths, heat shock at −20°C, followed by a 3h treatment with BSA 0.2%, 0.5% Triton X-100, 5% goat serum in PBS. Primary antibodies were incubated overnight at 4°C. Samples were washed three times and then incubated with secondary antibody mouse IgG_1_ Alexa Fluor® 488 overnight and mouse IgM Alexa Fluor® 4594 at 4°C. The following day, samples were washed three times with PBS 1X and incubated with Alexa Fluor™ 633-conjugated phalloidin for 2h. Transfected skin explants were analyzed by immunohistochemistry following the same methodology as whole embryos. anti-MelCAM mAb (sc18837, Santa Cruz Biotechnology) was incubated overnight at 4°C and evidenced by Alexa fluor® 488 and staining with solution Hoechst 33342 (2μM) (Thermo Scientific™) for two hours. Embryos and skin explants were imaged under AxioImager M2 (Apotome ZEISS), Confocal Leica SP8 microscope, and rendered with the Imaris Viewer 9.7.0. software.

*In vitro* melanocyte culture was analyzed by immunocytochemistry using anti-MelCAM mAb (Santa Cruz Biotechnology) and Phospho-Paxillin (Tyr31) polyclonal antibody (pAb) (ThermoFisher). Briefly, cells were fixed with 4% PFA for 20min. Cells were washed three times in PBS containing 0.1% Triton X-100 (PBST) and incubated with primary antibodies overnight at 4°C. Incubation with fluorophore-conjugated secondary Abs, mouse IgG_1_ Alexa Fluor® 488, and rabbit IgG Alexa fluor® 594 was carried out overnight at 4°C. For actin filament analysis, fixed cells were incubated with Alexa Fluor™ 633-conjugated phalloidin (Invitrogen™). The following day, cells were washed three times and incubated with Alexa Fluor™ 633-conjugated phalloidin for 1h, and slides were mounted with Fluoromount-G^TM^ with DAPI (Invitrogen). Immunohistochemistry on sections was imaged under a fluorescent microscope (Leica) equipped with a digital camera C10600 Hamamatsu Orca-R2, Germany, and a Confocal Leica SP8 microscope. Digitized images were adjusted in contrast and brightness using the Adobe Photoshop software.

### Imaging and image processing

Embryos were imaged under a stereomicroscope Leica Wild M10 equipped with a digital camera micropublisher 3.3 RTV and digitalization using Q-capture 6.0 software. Whole-mount staining was imaged under a Leica MZFLIII stereo zoom microscope and AxioImager M2 (Apotome) (ZEISS). Images were acquired using Leica Epifluorescence DM100, equipped with a digital camera HAMAMATSU ORCA-R^2^ C10600, and a Confocal Leica SP8 microscope. Skin explants images were taken as tiled z-stacks and were stitched together using ImageJ 1.0, and 3D rendering images were created using ImarisViewer 9.7.0. software. Three samples were analyzed by experimental conditions. All images were prepared using Adobe Photoshop 23.5.4 and Graphical abstract in Adobe Illustrator 27.2.

### Quantification and Statistical analysis

Images were analyzed using ImageJ 1.0, cell count with the cell counter plugin, and vessel length density with the Vessel Analysis plugin. Quantifying the vessel length was challenging to control due to the hyperplastic feather, so we divided each image into five sections. We measured vessel density for each section in the control and experimental series. Feather and filopodia length was measured by setting scale; then, the Straight Line tool was applied to measure the respective lengths. Paxillin analysis was identified based on the location within the cell compartment (nucleus, soma, or cell extension). All data plots and statistical analysis were determined by GraphPad Prism 9.4.1 software. Quantification results were compared using one-way ANOVA Dunnett’s multiple comparisons tests. A p-value of <0.05 was considered significantly different.

### RNA-seq data processing: Bulk RNA sequencing

We analyzed 3 biological samples per triplicate per condition for RNA sequencing: cranial and trunk skin samples (2 replicates on the cranial skin rescue *dsFBXO32+hFBXO32^WT^*). RNA extraction was performed using TRI reagent^R^ (Sigma-Aldrich) according to the manufacturer’s protocol. Briefly, dissected skin tissues were homogenized (either directly or after congelation at −80°C) in TRI reagent^R^ supplemented with 0,2 ml of chloroform for 1 ml of TRI reagent^R^. After centrifugation, the aqueous phase was transferred into a new tube and precipitated with 2-propanol. RNA pellets were washed with 75% ethanol and, after centrifugation and air drying, were dissolved in DEPC-H2O (SIGMA). RNA concentration was measured on NanoDrop^TM^2000c (ThermoFisher) and adjusted to 200 ng/μl for each sample.

The Novogene company performed cDNA synthesis and library preparation. Briefly, the first cDNA strand synthesis was synthesized randomly by hexamer primer M-MulV reverse transcriptase, and the second cDNA strand was synthesized using DNA polymerase I and RNAse. Libraries fragments were purified and sequenced on the Illumina Novaseq platform, and 150 bp paired ends reads were generated. The library was checked with Qubit, real-time PCR for quantification, and a bioanalyzer for size distribution detection. Quantified libraries will be pooled and sequenced on Illumina platforms according to effective library concentration and data amount. The index-coded samples were clustered according to the manufacturer’s instructions. After cluster generation, the library preparations were sequenced on an Illumina platform, and paired-end reads were generated. Raw data (raw reads) of fastq format were first processed through in-house perl scripts. In this step, clean data (clean reads) were obtained by removing reads containing adapter, reads containing ploy-N, and low-quality reads from raw data. At the same time, Q20, Q30, and GC content, the clean data, were calculated. All the downstream analyses were based on clean data with high quality.

Gene model annotation files were downloaded from the genome website directly, and we used them as reference genome *Gallus* gallus (ID: ensembl_gallus_gallus_grcg6a_gca_000002315_5). The index of the reference genome was built using Hisat2 v2.0.5, and paired-end clean reads were aligned to the reference genome using Hisat2 v2.0.5

### Quantification of gene expression level

Feature counts v10.5-p3 were used to count the read numbers mapped to each gene. Gene expression level quantification was assessed with the FPKM of each gene calculated based on the length of the gene and the reads count mapped of this gene. FPKM, the expected number of Fragments Per Kilobase of transcript sequence per Millions of base pairs sequenced, considers the effect of sequencing depth and gene length for the reads count at the same time and is currently the most used method for estimating gene expression levels.

### Differential expression analysis

Differential expression analysis of two conditions/groups (two biological replicates per condition) was performed using the DESeq2R 1.20.0 package. DESeq2 provides statistical routines for determining differential expression in digital gene expression data using a model based on the negative binomial distribution. The resulting p-values were adjusted using Benjamini and Hochberg’s approach for controlling the false discovery rate. Genes with an adjusted p-value≤0.05 found by DESeq2 were assigned as differentially expressed. Before differential gene expression analysis, the edgeR program package adjusted the read counts through one scaling normalized factor for each sequenced library. Differential expression analysis of two conditions was performed using the edgeR R 3.22.5 package (3.22.5). The p-values were adjusted using the Benjamini and Hochberg method. A corrected p-value of 0.05 and an absolute foldchange of 2 were set as the threshold for significantly differential expression.

### GO enrichment analysis of differentially expressed genes

GO enrichment analysis of differentially expressed genes was implemented by the cluster Profiler R package between *dsFBXO32+hFBXO32^WT^* vs *dsFBXO32+hFBXO32^G139S^*in cranial and trunk skin samples. GO terms with corrected p-value less than 0.05 were considered significantly enriched by differentially expressed genes.

### Gene Set Enrichment Analysis (GSEA)

GSEA is a computational approach to determine if a predefined Gene Set can show a significant, consistent difference between two biological states. The genes were ranked according to the degree of differential expression in the two samples, and then the predefined Gene Set were tested to see if they were enriched at the top or bottom of the list. Gene set enrichment analysis can include subtle expression changes. We use the local version of the GSEA analysis tool http://www.broadinstitute.org/gsea/index.jsp, GO, was used for GSEA independently.

### RNAseq data visualization

Volcano plots and bar charts for differential gene expression were generated in GraphPad 9.4.1, and Scatter plots using the online Plotly Chart studio.

## SUPPLEMENTARY FIGURE LEGENDS

**Figure S1:**
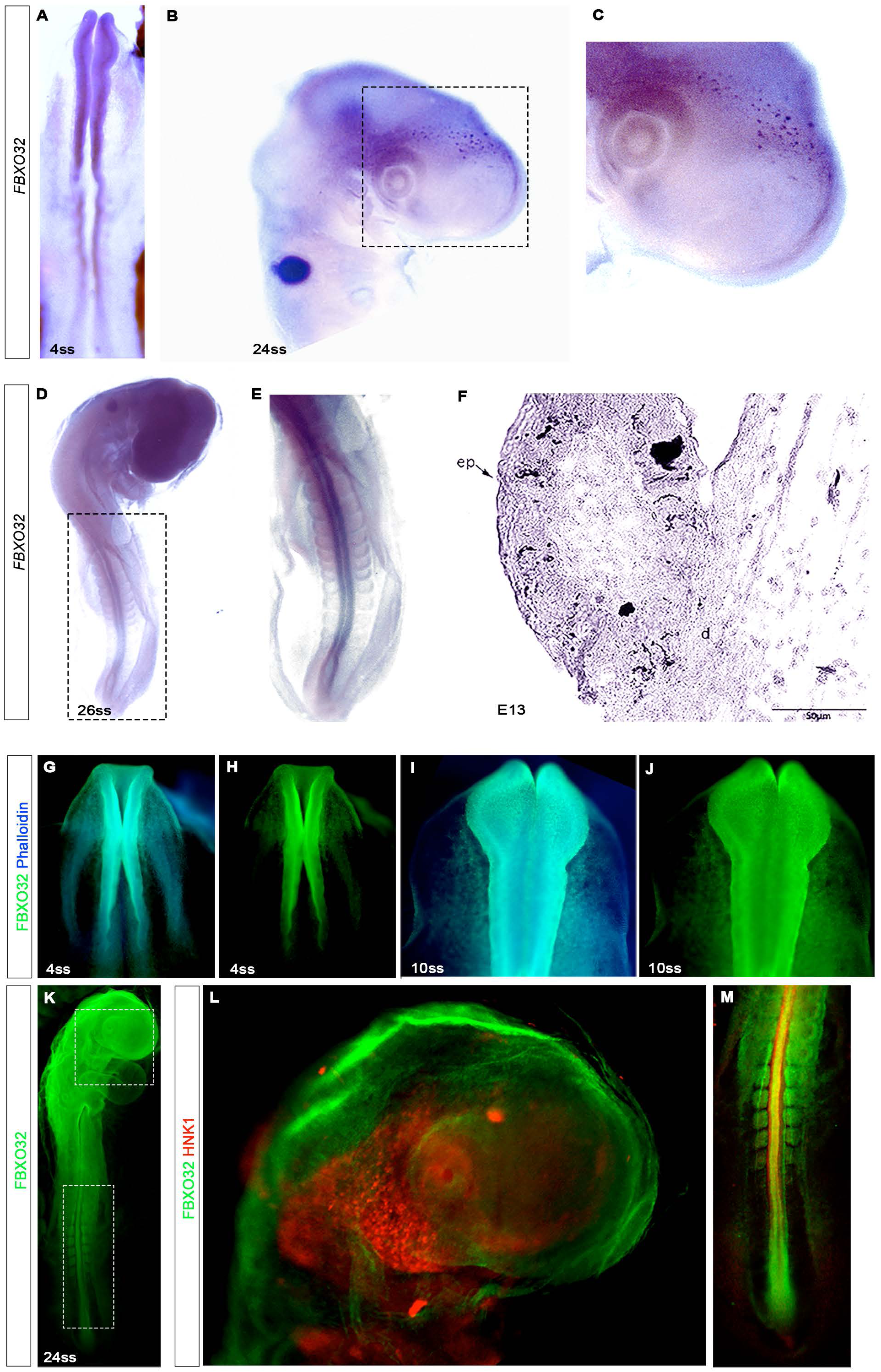
*FBXO32* is expressed in the CNC and TNC at different developmental stages. **A-F)** Whole mount *in situ* hybridization of *FBXO32* in the chick embryo **A)** At stage 4ss, expression of FBXO32 transcripts is detected in migratory NC. **B)** At stage 24ss, the expression of *FBXO32* transcripts is detected in the migrating NCC at the nasofrontal, retro-ocular, olfactory placode, and branchial-arches regions. **C)** Box area shows *FBXO32* transcripts in the migrating NCCs at the nasofrontal region. **D)** At stage 26ss, *FBXO32* transcripts are found in the developing head and the trunk. **E)** Box area shows *FBXO32* transcripts in the neural tube and the somites boundaries at the trunk. **F)** In a transversal feather section of embryo E13, *FBXO32* transcripts are localized in the melanocyte lineage at the epidermis layer. **G)** Immunolabelling of 4ss control embryo with anti-FBXO32 mAb (green) and phalloidin staining (cyan) shows colocalization of anti-FBXO32 mAb and phalloidin staining at the CNC. **H)** Control 4ss-embryo shows the accumulation of anti-FBXO32 mAb at the CNC region. **I)** Immunolabelling of 10ss control embryo shows colocalization of anti-FBXO32 mAb (green) and phalloidin staining (cyan) at the CNC. **K-M)** Immunolabelling of 24ss-TNC control embryo with anti-FBXO32 mAb (green) and anti-HNK1 mAb (red). **L-M)** Box areas showing delaminating NCCs colocalized with anti-FBXO32 mAb (green) and anti-HNK1 mAb (red) at the CNC and TNC, respectively.

**Figure S2:**
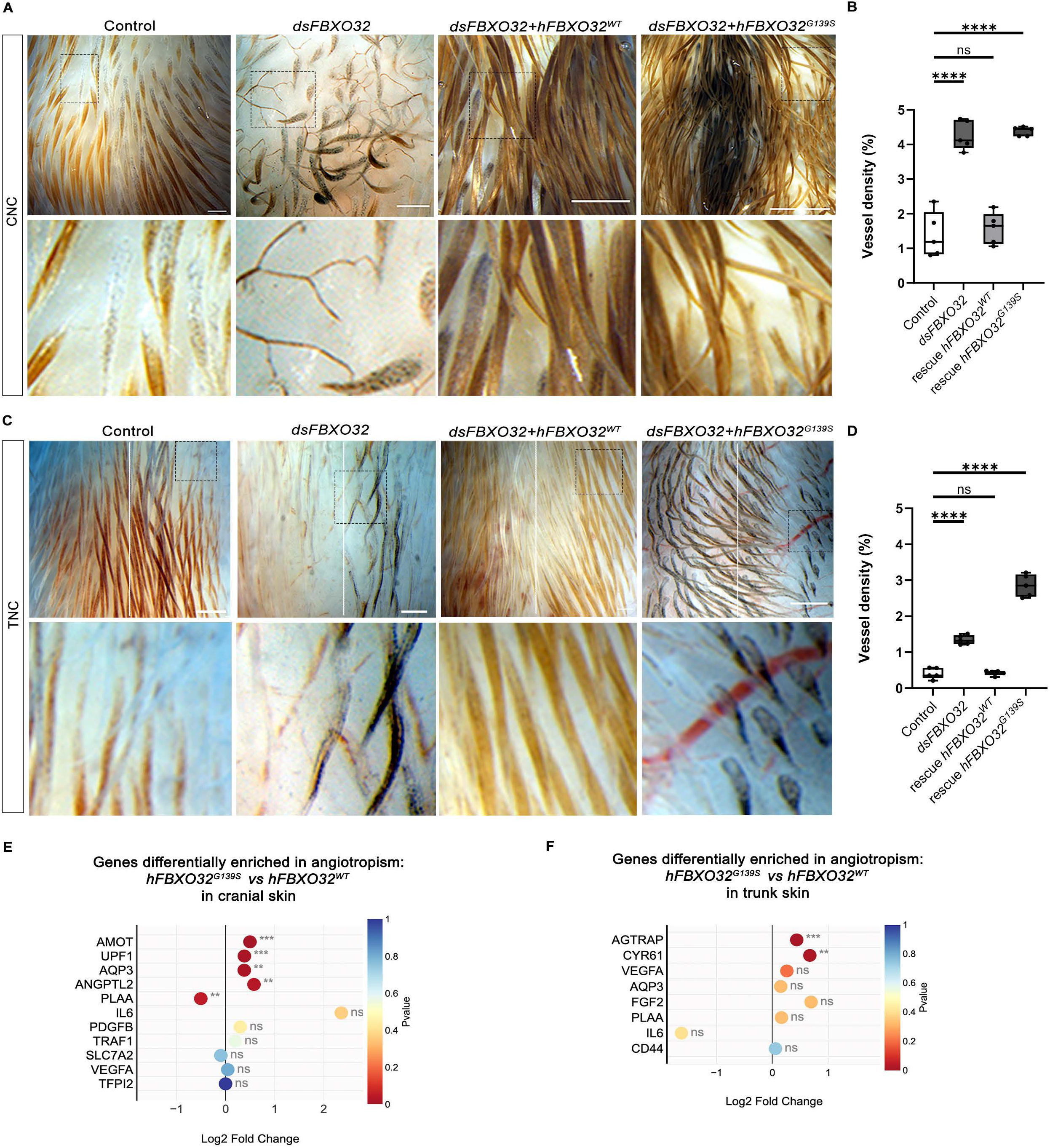
*FBXO32* enhances blood vessel density at the dermis. **A)** Dorsal views of E13 control embryos subjected to *FBXO32* silencing, *dsFBXO32+hFBXO32^WT^*, and *dsFBXO32*+*hFBXO32^G139S^* transfection in CNC. Blood vessel density increase was not evident in control and *dsFBXO32+hFBXO32^WT^*embryos. In embryos subjected to *FBXO32* silencing and *dsFBXO32*+*hFBXO32^G139S^*shown, blood vessel density increased at the dermis. Square boxes show the magnification of dorsal CNC regions. **B)** Box and whiskers plots comparing blood vessel density in the dorsal region of CNC embryos. **C)** Dorsal views of E13 control embryos, subjected to *FBXO32* silencing and co-electroporated with *dsFBXO32+hFBXO32^WT^* and *dsFBXO32+hFBXO32^G139S^* in TNC. Embryos subjected to *FBXO32* silencing and co-electroporated with *hFBXO32^G139S^* showed blood vessel density increased at the dermis. Square boxes show the magnification of dorsal TNC regions. **D)** Box and whiskers plots comparing blood vessel density in the dorsal region of TNC embryos. **E-F)** Scatter plot summarizing gene enrichment in angiotropism related genes in *dsFBXO32*+*hFBXO3^WT^-* vs *dsFBXO32*+*hFBXO32^G139S^*-treated cranial and trunk skins. The circle color indicates the significant level with the p-value<0.05. Vessel density was quantified with Fiji using a vessel analysis plugin. Each dot of the box and whiskers plots represents a mean±SD of an image divided into five segments. Bars indicate the mean; error bars, SD; one-way ANOVA with Dunnett’s multiple comparison test (\*\**P*<0.05, \*\*\*\**P*<0.0001, ns: not significant). Scale bar: 50µm

**Figure S3:**
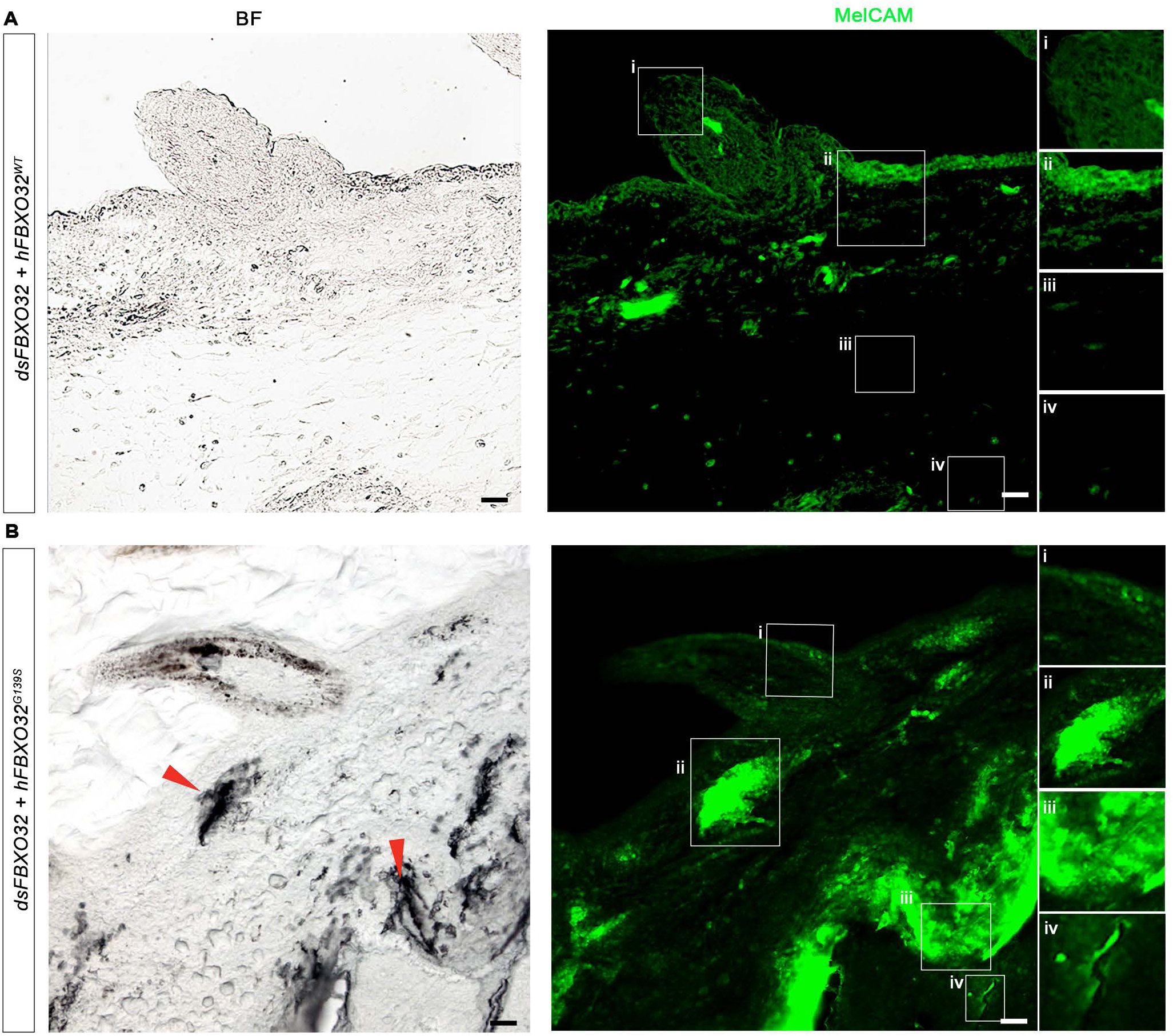
*FBXO32^G139S^*promotes the dissemination of melanocytes towards the dermis, Schwann cell, and vasculature in the skin. **A)** Bright-field image of a skin section of an embryo subjected to *dsFBXO32*+*hFBXO32^WT^* transfection in CNC showing differentiated melanocytes at the epidermis (left panel). On the right, immunostaining of the skin section in A) against anti-MelCAM mAb (green). Close-up views show slightly Melcam accumulation restricted to the epidermis in *i)* and *ii)*. **B)** Bright-field image of a skin section of an embryo subjected to *dsFBXO32*+*hFBXO32^G139S^*in the TNC (left panel). Red arrowheads highlight the aberrant accumulation of black patches of eumelanin in the dermis and Schwann cell compartments. Close-up view of the skin section in B) shows aberrant MelCAM mAb immunoreactivity (green) in the *i)* epidermis, *ii)* dermis, *iii)* Schwann deposit cells, and *iv)* perivascular deposit cells. Images correspond to optical sections. Scale bar: 50μm, and 25μm, respectively.

**Figure S4:**
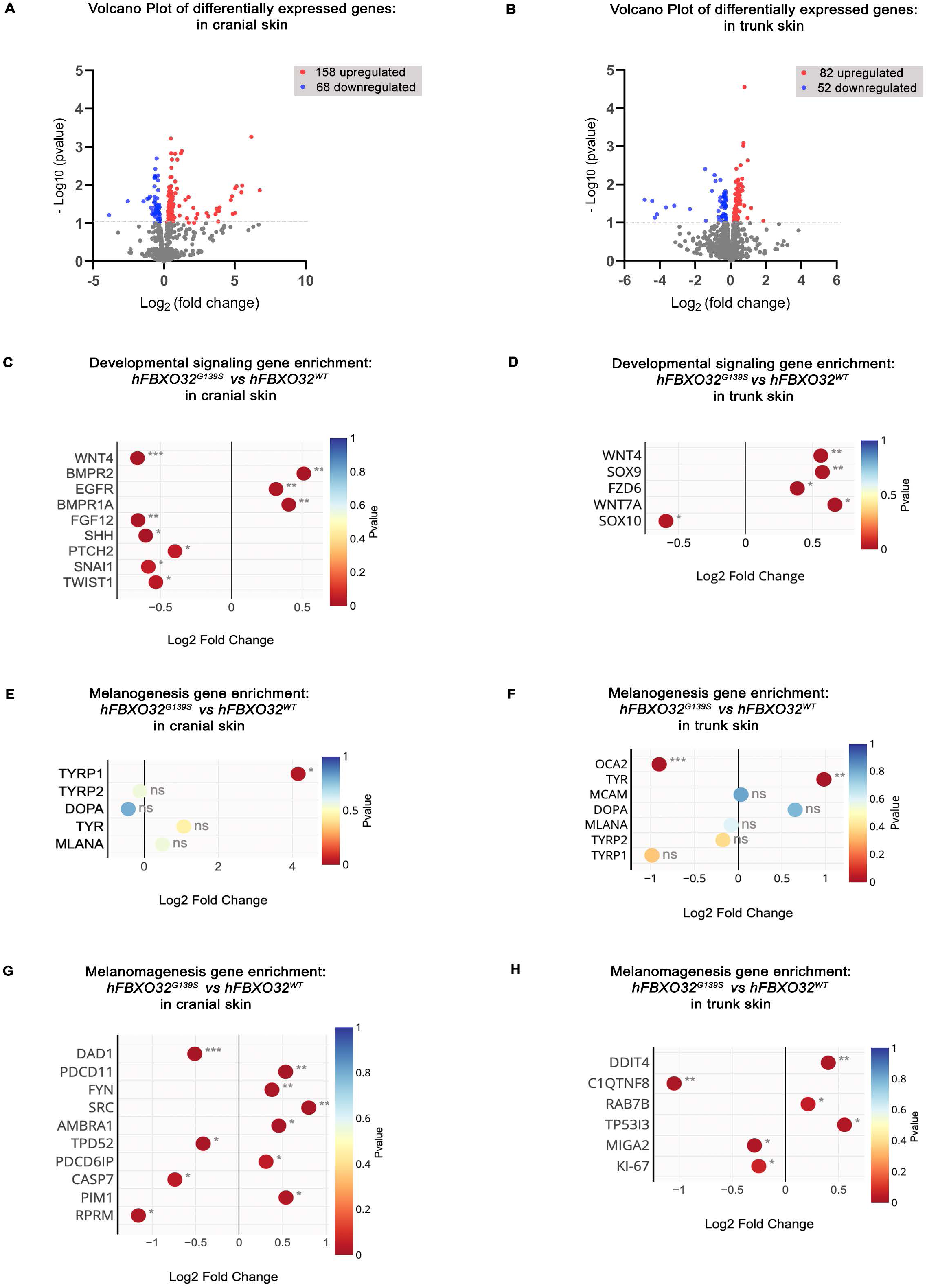
Differential expression of genes involved in developmental signaling, melanogenesis, and melanomagenesis in cranial and trunk skin. **A)** Volcano plot of genes statistically enriched or reduced in E13 cranial skin samples. **B)** Volcano plot of genes statistically enriched or reduced in E13 trunk skin samples. Red dots represent genes expressed at higher levels, while blue dots represent genes with lower expression levels. Y-axis denotes − Log_10_ (p-value), and X-axis denotes Log_2_ (fold change) values. The volcano plot was generated using GraphPad Prism9.5.0 software. The limit of greys dots spots delimitates the statistical significance set at log (1) (p<0.05). Scatters plots summarizing gene enrichment in C-D) developmental signaling, **E-F)** melanogenesis, and **G-H)** melanomagenesis, in between *dsFBXO32*+*hFBXO32^G139S^*- and *dsFBXO32*+*hFBXO32^WT^*-treated cranial and trunk skins, respectively. The circle color indicates the significant level with the p-value <0.05. Asterisks indicate significance: ∗*P*<0.05, ∗∗∗*P*<0.001. The scatter plots for gene expression were generated using Online Graph Marker-Ploty Chart Studio.

**Figure S5:**
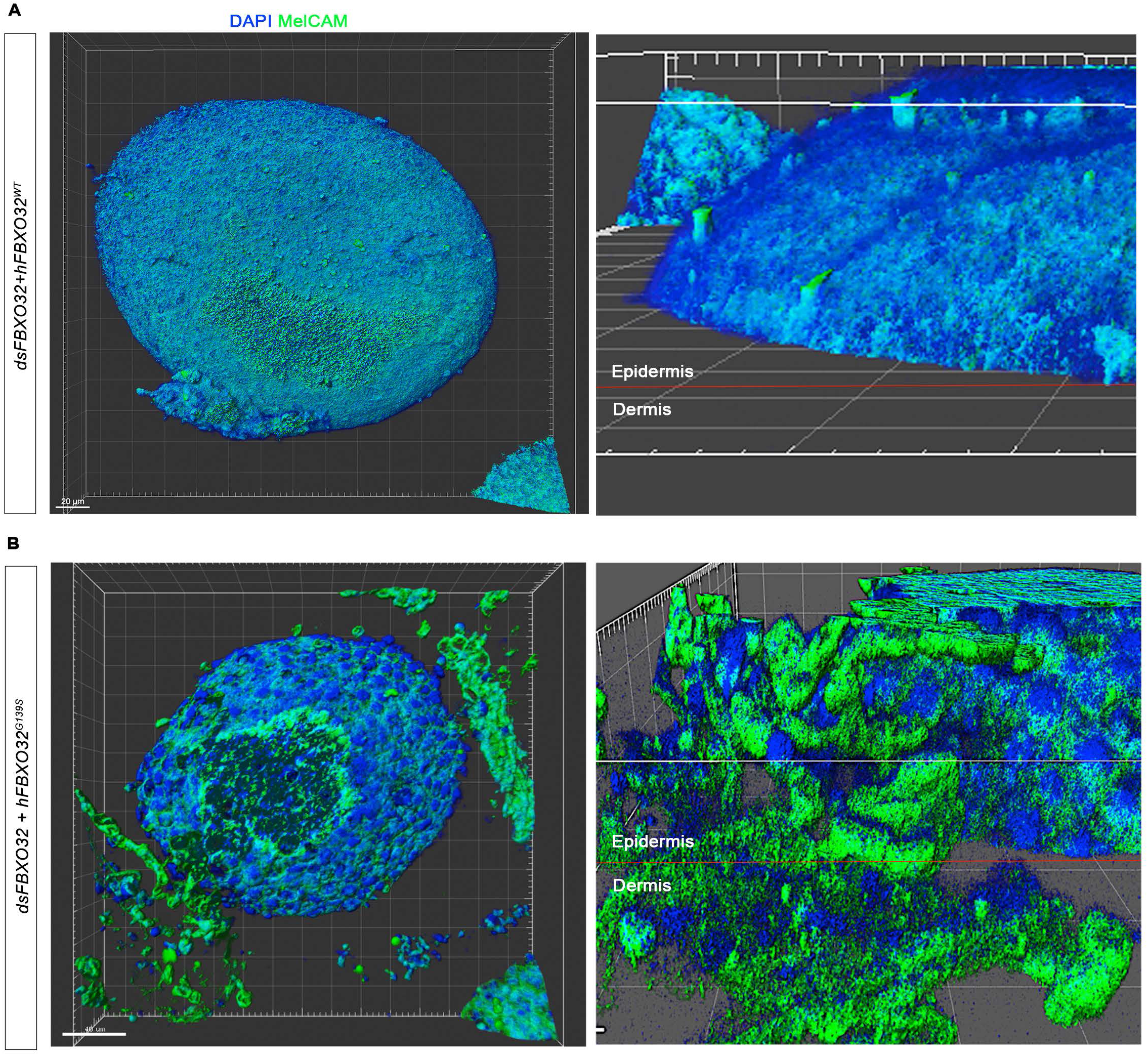
*hFBXO32^G139S^* induces melanocyte migration toward the dermis. **A-B)** 3D reconstructions of skin explants co-electroporated with *dsFBXO32+hFBXO32^WT^* and *dsFBXO32+hFBXO32^G139S^*showing immunoreactivity for anti-MelCAM mAb (green) and DAPI staining (blue) to assess melanocyte transformation. **A)** Close-up views showing MelCAM mAb immunoreactivity (green) in melanocyte filopodia extending toward the skin surface in the epidermis (right). **B)** Close-up views showing MelCAM mAb immunoreactivity (green) in melanocyte aggregates invading the dermis and accumulating in tissue disorganization (right). 3D rendering and visualization were created in IMARIS software of Z-stacks confocal images. The horizontal red dashed line indicates the boundaries between the epidermis and dermis layers. Scale bar: 20μm and 40μm, respectively.

**Figure S6:**
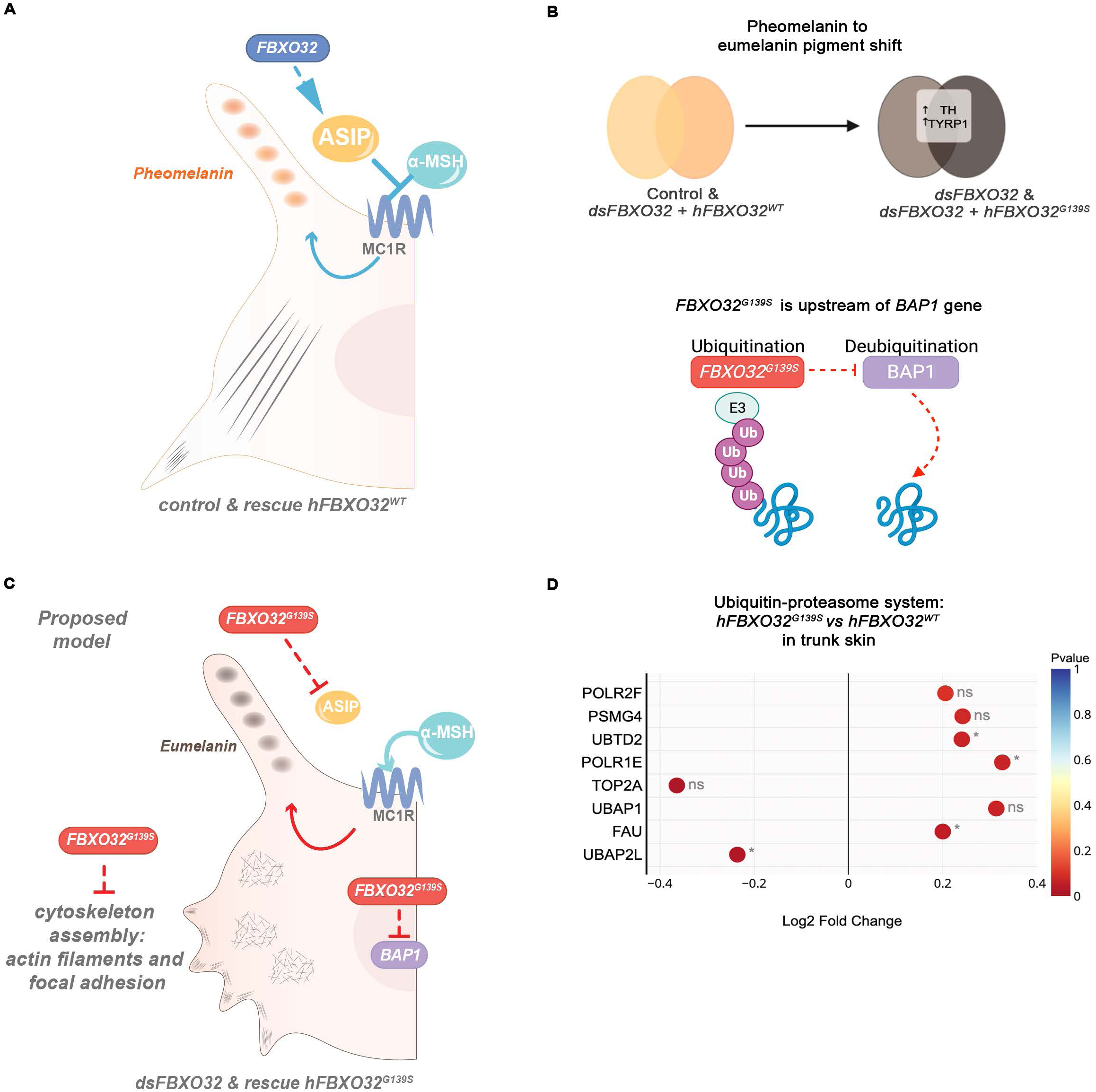
Schematic representation of key findings and proposal model. **A)** Schematic representation of control melanocyte illustrating the melanogenesis biosynthesis pathway, where *ASIP* prevents the binding of αMHS to MC1R to stimulate the synthesis of pheomelanin. Normal melanocytes exhibit long filopodia extensions with polymerized branched actin filaments and focal adhesion. **B)** When subjected to *dsFBXO32* silencing and *dsFBXO32*+*hFBXO32^G139S^* transfection, both cranial and trunk samples exhibit pheomelanin to eumelanin pigmentation shift and *BAP1* downregulation. **C)** We postulate that 1) *FBXO32^G139S^* is upstream of the *ASIP* gene, and its downregulation results in the binding of α-MSH to MC1R receptor, stimulating the synthesis of eumelanin, 2) *FBXO32^G139S^* impairs cytoskeleton architecture resulting in disassembly of focal adhesions and actin filaments, 3) *FBXO32^G139S^* downregulates *BAP1* gene in NCC-derived cells. **D)** Scatter plot summarizing gene enrichment in the ubiquitin-proteasome system between *dsFBXO32*+*hFBXO32^G139S^*and *dsFBXO32*+*hFBXO32^WT^*-transfected in cranial skin and trunk skins samples. This result highlights the impact of *FBXO32^G139S^* on ubiquitination genes.

